# The Fat/Hippo pathway drives photoperiod-induced wing length polyphenism

**DOI:** 10.1101/2024.04.05.588279

**Authors:** Erik Gudmunds, Aleix Palahí, David Armisén, Douglas G. Scofield, Freya Darling Eriksen, Abderrahman Khila, Arild Husby

## Abstract

Identifying the genetic mechanisms that translate information from the environment into developmental programs to control size, shape and color are important for gaining insights into adaptation to changing environments. Insect polyphenisms provide good models to study such mechanisms because environmental factors are the main source of trait variation. Here we studied the genetic mechanism that controls photoperiod-induced wing length polyphenism in the water strider *Gerris buenoi*. By sequencing RNA sampled from wing buds across developmental stages under different photoperiodic conditions known to trigger alternative wing developmental trajectories, we found that differences in transcriptional activity arose primarily in the late 5^th^ instar stage. Among the differentially expressed genes, the Fat/Hippo and ecdysone signaling pathways, both putative growth regulatory mechanisms showed significant enrichment. We used RNA interference against the differentially expressed genes Fat, Dachsous and Yorkie to assess whether they play a causative role in photoperiod induced wing length variation in *Gerris buenoi*. Our results show that the conserved Fat/Hippo pathway is a key regulatory network involved in the control of wing polyphenism in this species. This study provides an important basis for future comparative studies on the evolution of wing polyphenism and significantly deepens our understanding of the genetic regulation of insect polyphenisms.

## Introduction

The size and shape of insect wings are determined by the action of developmental genetic programs in conjunction with hormonal cues that adjust growth according to endogenous and exogenous environment (Tripathi and Irvine, 2022). Wing size has an environmentally-sensitive component in many insect species to allow proper scaling of wing size to body size (Nijhout and Callier, 2015), and this relationship is impacted by a number of environmental factors such as temperature and diet (Bakker, 1959; Robertson, 1963; Atkinson, 1994; Partridge *et al*., 1994; Nijhout, 2003b). For some species environmental factors can induce discrete wing length variation, termed wing polyphenism (Hayes *et al*., 2019). Wing polyphenisms are generally considered as examples of adaptive plasticity that acts to increase the fitness of an individual in relation to selective agents that can be predicted from environmental cues (Nijhout, 2003a). A fundamental and largely unanswered question pertaining to wing polyphenism is how the conserved genetic and endocrine pathways governing wing development have evolved to generate discrete morphological variation based on input from environmental cues.

In *Drosophila*, organ intrinsic (e.g. morphogens) and extrinsic (hormones) factors act during development to regulate the final wing size of individuals (Tripathi and Irvine, 2022), where the plastic growth largely occurs through the endocrine axis of regulation (Mirth and Shingleton, 2019). In this system, insulin-like peptide (ILP) signaling canonically acts through the insulin/insulin-like growth factor signaling (IIS) and target of rapamycin (Tor) pathways to adjust size in accordance to nutrient levels (Brogiolo *et al*., 2001; Hietakangas and Cohen, 2009), whereas ecdysteroid signaling regulates both size and patterning (Gokhale *et al*., 2016; Nogueira Alves *et al*., 2022) by mediating the regulation of genes and pathways involved in wing morphogenesis (Herboso *et al*., 2015; Parker and Struhl, 2020; Strassburger *et al*., 2021; Perez-Mockus *et al*., 2023). Notably, the regulatory function of these two hormones in insect wing development is conserved over considerable evolutionary distances (Herboso *et al*., 2015; Nijhout and Callier, 2015; Nijhout, Laub and Grunert, 2018) and both have been shown to regulate morph determination in wing polyphenic species, as well as play a role in other polyphenisms (Rountree and Nijhout, 1995; Xu *et al*., 2015; Vellichirammal *et al*., 2017; Fawcett *et al*., 2018; Nijhout and McKenna, 2018; Smýkal *et al*., 2020; van der Burg *et al*., 2020). Hence, a likely commonality for polyphenisms is that they evolve through modification of the conserved signal transduction pathways for environmentally-sensitive growth and development.

It is well-established that the environmental cues driving polyphenisms are translated into the endocrine signaling environment of insects, which in turn directs the developmental trajectory of individuals (Nijhout, 2003a). Yet, how hormones act to regulate the proximate genetic pathways of growth and patterning in the focal tissues to generate alternative morphologies remains largely unclear (Gotoh *et al*., 2015). However, despite the general lack of genetic experimentation tools for species displaying alternative morphologies, some recent studies have unraveled a molecular switch system operated by the expression pattern of insulin receptor (InR) paralogs in the brown planthopper *Nilparvata lugens* (Xu *et al*., 2015; Lin *et al*., 2016). Here, growth induced by the IIS pathway is tuned down when an alternative InR (InR2) is expressed, likely due to the formation of an InR1/InR2 receptor dimer that is less capable of signal transduction than the canonical InR1/InR1 dimer (Xu *et al*., 2015).

Despite the fact that a role for the IIS pathway has also been established in other hemipteran species (Fawcett *et al*., 2018; Smýkal *et al*., 2020) – which suggests a conserved role in the regulation of wing dimorphisms – we recently showed that the IIS pathway does not control photoperiodically-induced wing polyphenism in the water strider *Gerris buenoi* (Gudmunds *et al*., 2022). Interestingly, nutrition does not act as a cue for the induction of alternative wing morphs in *G. buenoi* (Gudmunds *et al*., 2022), in contrast with the nutritionally-sensitive and IIS-regulated polyphenisms in the brown planthopper and the soapberry bug (*Jadera haemotoloma*, Fawcett *et al*., 2018; Lin *et al*., 2018). This interspecific variation in inductive cue and pathway utilization in hemipteran wing polyphenism thus poses the question of whether the evolution of polyphenisms is constrained to specific genetic routes as a consequence of the identity of both the selective agent and the triggering environmental cue. Given that the only hemipteran wing polyphenisms for which specific genetic pathways have been identified to date are regulated through nutrition, the exploration of the evolution of wing polyphenism necessitates studies in species where nutrition is not the morph-determining cue.

In this study, we used RNA sequencing (RNA-seq) and RNA interference (RNAi) to explore the mechanisms underlying seasonal wing length variation in the water strider *Gerris buenoi*. Wing length in this species is strongly regulated by photoperiod, and shows the clear bimodal distribution characteristic of a polyphenism both in the wild (Spence, 1989) and in the lab (Gudmunds *et al*., 2022). When reared in 12L:12D nearly 100% of individuals develop into the long-winged (macropterous) morph and close to 100 % of individuals reared in 18L:6D develop into the short-winged (micropterous) morph. The strong correlation between photoperiod and adult wing morph in *G. buenoi* greatly facilitates transcriptome sampling throughout development, as photoperiod provides a strong predictive basis of the developmental trajectory i.e. which adult wing morph the individual will develop into. To our knowledge, this is the first tissue-specific gene expression study of wing polyphenism in direct association to the environmental cue that operates the developmental switch (but see Vellichirammal, Madayiputhiya and Brisson, 2016; Zhang *et al*., 2021 for alternative approaches). Our results yield new insights into the genetic basis of wing polyphenism and support the idea that distinct genetic pathways underlying wing polyphenism in hemipterans have been coopted depending on the environmental cue.

## Material and Methods

### Water strider husbandry and photoperiod treatments

The *G. buenoi* population used in this study was originally collected in Toronto, Canada, during 2012 but has been replenished several times since with individuals from the same area. It has continuously been reared in the lab in photoperiod conditions that generated different wing morphs. The parental population for the experiments was reared at ∼20°C (room temperature, RT) in a 22:2 light:dark photoperiod. The replenishment of adults to the parental population occurred mainly from individuals that had been reared in 18L:6D or 12L:12D at 25°C, thus constituting a mix of macropterous and micropterous individuals (see Figure 1A). Feeding occurred five times a week with frozen crickets (*Acheta domestica*). All experiments with photoperiod occurred in growth rooms with ∼80 μEinstein (9400 lux) light intensity at 25°C constant temperature. Aeration with air stones connected to an air pump was used to ensure that the water surface was kept clean and remained suitable for water striders.

**Figure 1.**
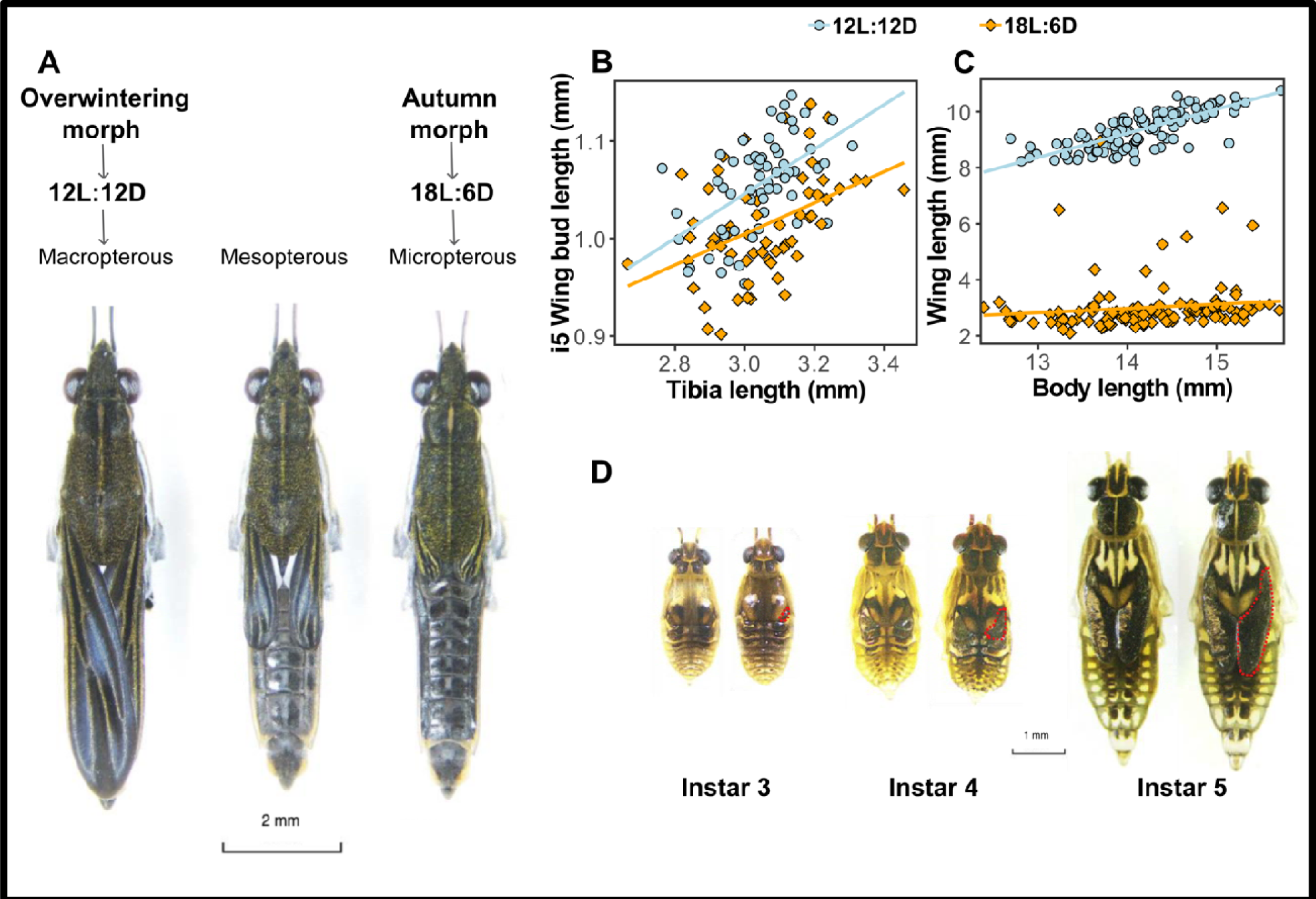
Development of *G. buenoi* as nymphs and upon reaching adulthood. **A**) From left to right, macropterous, mesopterous and macropterous male individuals. **B)** Correlation between i5 wing bud length and tibia length in individuals reared in either 12L:12D and 18L:6D. **C)** Correlation between adult wing length and body length in individuals reared in either 12L:12D or 18L:6D. **D)** Left to right, instar three, four and five individual pairs. Within each pair, the left individual was reared in 12L:12D and th right individual in 18L:6D. Wing buds are encircled with the red dotted line.

### Size measurements

The data on adult body and wing length were taken from previously published material (Gudmunds *et al*., 2022). Here, individuals were reared in different nutrient regimes in either 12L:12D or 18L:6D until adulthood and then body length and wing length were measured using ImageJ (version 1.53 k). The data includes both males and females. Measurement of instar five (i5) wing bud and tibia length were performed by collecting and taking photos of i5 exuvia (Nikon SMZ800 stereo microscope with Nikon DS Fi1 camera).

### Experimental set-up and sampling for RNA sequencing

With the intention to characterize gene expression changes mediating the plastic response to photoperiod, individuals were reared in both short (12L:12D; generating ∼100% macropterous individuals) and long day conditions (18L:6D; generating ∼100% micropterous individuals). Nymphs from both photoperiods were sampled at different time intervals after eclosion into both instar four (i4) and i5 to use for RNA extractions and sequencing.

These nymphal stages were chosen based on a previous experiment where we showed i4 to concentrate most change in sensitivity to photoperiod, and i5 to be when the adult wing tissue is specified and undergoes differential growth (Gudmunds *et al*., 2022). We firstly assessed the developmental duration of instar three (i3), i4 and i5 in 12L:12D and 18L:6D at 25 °C constant temperature to ensure that the sampling of individuals between the two photoperiods represented proportional development within each instar (Supplemental Table S1). For that purpose, we collected eggs randomly from the stock population and reared individuals in groups in either photoperiod. Upon moulting into i3, we isolated individuals in plastic cups, where they were fed once a day with a single cricket, and monitored moulting events every day at mid-photophase. The developmental duration (recorded in days) was analyzed using a two-sided Student’s t-test. For each photoperiod, we started the experiment with 60 individuals but due to mortality the final sample size was 55 for 12L:12D and 49 for 18L:6D. The outcome of the developmental duration experiment led us to sample individuals at the following four sampling timepoints (relative to total instar duration): i4 30% (hereafter instar four early; i4E), i4 ∼85% (instar four late; i4L), i5 ∼19% (instar five early; i5E) and i5 ∼67% (instar five late; i5L). In chronological time, we sampled at i4 24 hours after eclosion (hae) for i4E, i4 72 hae for i4L, i5 24 hae for i5E, and i5 72 hae in 18L:6D (corresponding to 64% of development) and 96 hae in 12L:12D (corresponding to 69% of development) for i5L, since i5 duration in 12L:12D is on average one day longer than in 18L:6D (Supplemental Table S1).

This setup was used to sample males for RNA extractions to decrease variation in developmental stage when sampling. For that purpose, we collected eggs from the stock population and randomly distributed them into two growth rooms with either 12L:12D or 18L:6D. Hatched individuals were transferred to a growth box with *ad libitum* food until they reached i3, at which point they were transferred to another growth box. This box was carefully monitored each day for i4 eclosion events that occurred within a time window of ∼8 hours (midday ± 4 hours). Individuals that molted into i4 during this window were isolated and then sampled at the different developmental timepoints of interest (see above). If an i4 eclosion event occurred outside of the sampling window, the individual was kept in the box so that it could be sampled for the i5 timepoints, for which the procedure was repeated. Nymphal density in all stages after the hatching box was always <30 individuals/box, to ensure optimal growing conditions and to minimize the appearance of macropterous individuals in 18L:6D, which can be induced by high rearing densities (Gudmunds *et al*., 2022). A subset of individuals was reared until adulthood to score wing morph and verify the expected frequencies. This control confirmed that 12L:12D induces macropterous adults (100%) and that individuals reared in 18L:6D mostly become micropterous (∼90 %). The higher proportion of macropterous individuals obtained in 18L:6D compared to previous published data (Gudmunds *et al*., 2022) from the same photoperiod was likely due to a higher than expected effect of density in the growth boxes. To avoid any bias due to the time of the day when sampling, all individuals were sampled at midday in each photoperiod, six and nine hours after light on in 12L:12D and 18L:6D, respectively. Individuals for RNA extraction (males) were picked with ethanol-wiped forceps and immediately transferred into tubes with RNAlater (Invitrogen) and 1% Tween20. The tubes were then stored at 4°C for 2 hours and later at –80°C until further processing.

### Dissection and RNA extraction

The tubes with individuals were thawed on ice before dissection in ice-cold PBS with 1% Tween 20. With fine forceps, the left and right fore-wing buds were removed and transferred into 20 μl Trizol (Invitrogen). This procedure was repeated until a pool of either five (for i4E and i4L) or three (for i5E and i5L) pairs of wing buds had been collected. The wing buds were then homogenized with the use of a dissection needle, and 480 μl of Trizol was added and the tubes were thoroughly vortexed. The samples were stored at –20°C until further processing, which occurred when all dissections for all timepoints had been completed.

RNA was extracted following the standard Trizol protocol with a few modifications. First, the Trizol tubes were thawed and vortexed, and 100 μl of 24:1 chloroform-isoamyl alcohol was added prior to hand-shaking for 15 seconds and 3-minute incubation on ice. The mixture was then centrifuged at 12000g and 4°C for 15 minutes, and the supernatant transferred to a new tube. 250 μl ice-cold isopropanol and 1 μl glycogen (20 mg/ml) were added followed by overnight incubation at –20°C. Precipitated RNA was collected by centrifuging for 30 minutes at 12000g at 4°C. After discarding the supernatant, the RNA pellet was washed two times with ice-cold 75% ethanol. After air-drying, 30 μl of water was used to re-suspend the RNA pellet and incubate it at 65°C for 5 minutes, prior to a treatment with DNase I (Thermo Fisher). To remove the DNase and the glycogen in the solution, the samples were purified with GeneJET RNA cleanup and concentration kit (Thermo Fisher) according to the manufacturer’s protocol. The final elution of RNA was done with 20 μl of water. Agilent Bioanalyzer (RNA Nano kit) was used to validate the RNA integrity and estimate the concentration.

### RNA sequencing, read processing and mapping

Library preparation and next generation sequencing was performed by National Genomics Infrastructure (NGI) in Stockholm. Firstly, 46 libraries (5 – 6 per combination of developmental stage and photoperiod) were prepared using the Illumina TruSeq RNA poly-A selection kit for each individual sample. Following, all pooled libraries were sequenced in a single Illumina NovaSeq6000 S4 lane with 150 base pair paired end reads. Following demultiplexing, sequencing adapter filtering was done with Cutadapt v2.3, with a minimum overlap length between adapter and read of 8 bp and a minimum read length of 20 bp (Martin, 2011). Trimmed pair-end reads were mapped to the *G. buenoi* reference genome (see below) with STAR v2.7.9a (Dobin *et al*., 2013), using the two-pass mode. The number of total mapped reads per library is detailed in Supplemental Table S2.

### Sample filtering and differential expression analysis

After mapping, read-pair raw counting was performed for all annotated genes using the *featureCounts* function as implemented in subread v2.0.0 (Liao, Smyth and Shi, 2014) with the – p argument. Analysis of the raw counts was performed using R v4.2.1 (R Core Team, 2022) package *edgeR* v3.38.4 (Robinson, McCarthy and Smyth, 2010). Normalization factors were calculated for each library using the trimmed mean of M-values (TMM) method, in order to compute the counts per million (CPM) normalized expression for each transcript. Low-expression transcripts (i.e., transcripts with five or less CPM in at least three of the samples) were removed from further analysis.

Principal Component Analysis (PCA) was performed with the log2-transformed CPM counts to assess the clustering of all libraries within the different treatment groups. Three libraries (i5E_18_3, i5L_18_2 and i5L_18_5) were discarded due to low total number of mapped reads (Supplemental Table S2). Based on the PCAs obtained for all libraries (Figure S1), library i5L_18_1 was removed due to poor clustering. PCAs obtained for stage-specific libraries (Figure S2B, D) supported removal of two more samples from further analysis (i5L_12_4 and i4L_18_1). Finally, sample i4L_12_3 was discarded based on the PCA plot with the thousand most expressed genes (Figure S3). The remaining 39 libraries (3 – 6 per stage and photoperiod) were used to fit a negative binomial generalized linear model, to test for significant differential expression (DE) within developmental stages across photoperiods with a quasi-likelihood F-test (QLF). The contrasts used were: 12L:12D i4E vs 18L:6D i4E, 12L:12D i4L vs 18L:6D i4L, 12L:12D i5E vs 18L:6D i5E and 12L:12D i5L vs 18L:6D i5L. P-values were adjusted for multiple testing with the false discovery rate (FDR) method, and all genes with FDR < 0.01 and |logFC| > 0 were defined as differentially expressed.

### Gene Ontology Enrichment Analysis

Gene ontology (GO) enrichment analysis was performed in R v4.2.1 package *topGO* v2.48.0 (Alexa and Rahnenfuhrer, 2023). The gene universe included all genes found in the reference genome that were functionally annotated with at least one GO term. Our list of genes of interest corresponded to the list of differentially expressed genes between photoperiods for the same developmental stage. GO terms were considered as significantly enriched if they presented a Classic score < 0.05 (i.e. Fisher exact tests for each GO term, comparing the expected number of differentially expressed genes at random with the observed number of significantly DE genes).

### Gene Correlation Network Analysis

Signed, weighted gene correlation networks were created with the R v4.2.1 package *WGCNA* v1.72 (Langfelder and Horvath, 2008). The normalized read counts after low-expression gene filtering were used as input for the analysis. The networks were built using the blockwiseModules() function using a soft-threshold power of 16. Modules were required to include a minimum of 30 genes, and the resulting modules with a cut height < 0.25 (equivalent to a correlation > 0.75) were merged. The gene clusters identified by *WGCNA* that showed the highest differences in eigengene correlation between i5L photoperiods (Figure 4A) served as genes of interest in GO enrichment analyses, using the same parameters as before.

### DNA sequencing and genome assembly

A male and a virgin female were collected from descendants of the inbred population established for the first *Gerris buenoi* genome sequencing (Armisén *et al*., 2018). They were crossed and both parents and eleven offspring were sent to Dovetail Genomics (CA, USA) for DNA extraction and sequencing with Hi-C/Hi-Rise libraries. Using the published *Gerris buenoi* genome within the i5k initiative (Poelchau *et al*., 2015; Armisén *et al*., 2018) as input assembly, two successive assembly rounds were performed. Following scaffolding, the resulting assembly was polished using five rounds of Pilon v1.24 (Walker *et al*., 2014). For each round, four sets of Illumina HiSeq 2000/2500 whole-genome DNA-Seq paired-end reads (SRA accessions SRR1197265, SRR1197266, SRR1197267, SRR1197268, providing a total coverage of ∼87x) were aligned to the genome sequence using the bwa v0.7.17 *mem* algorithm (Li, 2013), sorted using SAMtools v1.12 (Danecek *et al*., 2021), and polished with Pilon v1.24 in diploid mode. Completeness of the final polished assembly was assessed using sets of universal single-copy orthologs for Insecta from OrthoDB v10v1 (Kriventseva *et al*., 2019), as implemented in BUSCO v5.2.2 (Manni *et al*., 2021).

### Genome annotation

The repeat library was developed from the genome using tools within the Dfam Consortium TETools Docker image v1.4 (github.com/Dfam-consortium/TETools). *De novo* repeats were determined using RepeatModeler v2.0.2 (Flynn *et al*., 2020) while enabling its LTR discovery pipeline. Discovered repeats were combined with Hemiptera-level repeats available within RepeatMasker v4.1.2-p1 (Smit, Hubley and Green, 2013) and then screened to remove sequences that matched repeat-containing proteins from the 16 Hemiptera taxa found within OrthoDB v10v1 (Kriventseva *et al*., 2019) and proteins from the i5k *G. buenoi* v1.1 genome annotation using RMblastn v2.11.0+. The polished genome assembly was soft-masked for repeats in the screened library together within low-complexity repeats using RepeatMasker 4.1.2-p1.

The soft-masked genome was annotated using the BRAKER pipeline (Stanke *et al*., 2008; Hoff *et al*., 2016; Brůna *et al*., 2021), which separately examines RNA-Seq and protein evidence together with *de novo* gene discovery with GeneMark v4.72 (Brůna *et al*., 2021), and produces a final consensus annotation using TSEBRA v1.0.3 (Gabriel *et al*., 2021). RNA-Seq evidence was provided by Illumina paired-end and single-end RNA-seq. All RNA-seq reads were aligned to the genome using STAR 2.7.2b (Dobin *et al*., 2013) and provided as sorted BAMs to BRAKER. The protein evidence included sequences from the 16 Hemiptera taxa found within OrthoDB 10v1 (Kriventseva *et al*., 2019) together with sequences from the i5k *G. buenoi* 1.1 genome annotation. Functional annotations and GO terms were added to the consensus annotation by comparing predicted proteins to *Drosophila melanogaster* proteins from genome release 6.45 provided by FlyBase FB2022_02 (Gramates *et al*., 2022), and attaching information provided by the top hit as determined by blastp v2.12.0+ (Camacho *et al*., 2009). The annotation was further augmented by direct alignment of 1241 manually curated unique proteins for known genes derived from the i5k *G. buenoi* v1.1 genome and annotation. These were aligned against the polished genome assembly using Exonerate v2.4.0 (Slater and Birney, 2005) in protein2genome mode with minimum 90% identity. Approximately 89% of curated proteins had a single hit against the polished genome, and 1.9% found no hit. Of the remaining proteins with multiple hits, the best hit was kept, as well as duplicate hits with >98%. In total, 1214 gene models were constructed using the manually curated proteins, with functional annotations and GO terms added from FlyBase FB2022_02 (Gramates *et al*., 2022) as described above. These gene models were integrated into the consensus annotation after removing overlapping consensus gene models.

### RNA interference

Preparation of dsRNA was done as described in (Gudmunds *et al*., 2022). Briefly, the primers used for each gene are listed in Supplemental Table S3. I4 individuals were injected with 0.3-0.4 μl and i5 with 0.4-0.6 μl. The concentration of injected dsRNA was 5 μg/μl or higher for all genes except Yki which was injected at 1000 ng/μl to avoid excessive mortality. Both i4 and i5 individuals were injected approximately 24 hae into either instar. Individuals used for RNAi hatched in 12L:12D and were then reared at a moderate density until they were injected in instar four or five. After injection individuals were again reared in either 12L:12D at moderate densities. All individuals used for RNAi were fed with *Acheta domestica* juveniles five times per week and temperature was kept constantly at 25°C. Wing morphs were scored using previously described criteria (Gudmunds *et al*., 2022). Individuals showing severe wing shape abnormalities were not included in the scoring, whereas individuals with slight abnormalities like faint wing venation, small differences in size between the left and right wing, or slightly curved wings were included. The number of individuals surviving to adulthood and used in the wing morph frequency data is shown above each bar for each gene in Figure 5B.

Knockdown validation of dsRNA treatments producing significant differences in wing morph frequencies was carried out with reverse-transcriptase quantitative PCR (RT-qPCR). Here, total RNA was extracted using Trizol (Qiagen) from i5 males (6-8 biological replicates per treatment) sampled 2 days after dsRNA injection performed as above. The RNA was treated with DNase I (Thermo Scientific) and then re-purified before RNA integrity control on Agilent Bioanalyzer and cDNA synthesis (RevertAid First Strand cDNA synthesis kit, Thermo Scientific). 3 μg of RNA was used for cDNA synthesis and 1 μl of 1:5 diluted cDNA was used in the RT-qPCR reaction triplicate (Luminaris Color HiGreen qPCR master mix, Thermo Scientific). Efficiencies of primers was calculated from standard curves to ensure similar amplification efficiencies between quantified genes. The ribosomal protein S26 (RPS26) was used as reference gene as it was closest in cycle threshold values and primer efficiencies for experimental genes among in total three tested reference genes. The ddCt method was used to calculate fold differences in gene expression between treatments. Primer sequences are listed in Supplemental Table S3.

## Results

### Photoperiod and wing development in G. buenoi

*G. buenoi* develops through five nymphal instars before molting to an adult with fully patterned wings (Figure 1A). Wing development occurs within wing buds, which appear as morphologically distinct structures in i3 and subsequently undergo incremental changes in shape and size in each successive molt (Figure 1D). However, it is likely that the wing identity of the wing bud progenitor tissue is specified already during embryogenesis as in other Hemimetabolous insects (Fernandez-Nicolas *et al*., 2022; Ohde, Mito and Niimi, 2022). Post-embryonic wing development in water striders is variable, with some species developing distinctly different wing bud sizes depending on adult wing morph fate. For example, individuals destined to become macropterous can in some species display wing buds which are markedly larger than those destined to grow shorter wings (Andersen, 1982). Such differences are not visually apparent in *G. buenoi*, indicating that the growth differentiation generating the alternative wing morphologies occurs during i5. However, using digital methods we found that i5 wing buds of individuals reared in 12L:12D i.e., destined to become macropterous, are slightly longer (means: 12L:12D – 1.056 mm, 18L:6D – 1.014 mm, average difference = 4%, F_1,117_ = 19.9, *P* < 0.01) than those of individuals reared in 18L:6D i.e., destined to become predominantly micropterous. The differences in wing bud length are independent from body-size variation (F_1,117_ = 0.42, *P* = 0.52, Figure 1B). These results show that the photoperiod-responsiveness of wing development in *G. buenoi* starts in i4 at the latest, with the first visible effects on the wing buds appearing as a quantitative difference in i5. It is also apparent that the wing bud growth for both photoperiod regimes scales with body size (18L:6D: R^2^ = 0.19, F_1,58_ = 14, *P* < 0.01, 12L:12D: R^2^ = 0.29, F_1,57_ = 23, *P* < 0.01), in contrast to the wing morph-specific growth occurring during i5 (Figure 1B).

In adults, the wing size of individuals reared in 12L:12D positively scales with body size (R^2^ = 0.61, d.f = 1, F = 178, *P* < 0.01; Figure 1C), reflecting a necessity to match wing size to overall body size to enable proper flight capability, whereas the wing size of individuals reared in 18L:6D is unrelated to body size (R^2^ = 0.02, d.f. = 1, F = 1.83, *P* = 0.17; Figure 1C). Taken together, these results show that the relationship between body and wing tissue growth in 18L:6D markedly shifts after the molt to i5, when the mechanism underlying the wing polyphenism decouples the wing tissue from the whole-organism signals responsible for coordinating growth between body parts.

### Temporal effects of photoperiod on expression dynamics in wing progenitor tissue

To investigate the differences in gene expression underlying the photoperiod-induced wing morph, we extracted and sequenced RNA from wing buds of individuals reared in 12L:12D and 18L:6D at four different timepoints of development, two in i4 and two in i5. After filtering out seven libraries from various treatments (see Methods for details), principal component analysis (PCA) showed evidence of a strong clustering of libraries prepared from tissue from the same developmental timepoint and photoperiod. This is a clear indication that the staging effort during sampling was effective (Figure 2A). The obtained clustering patterns are also indicators of significant gene expression dynamics across development. When comparing the same developmental stage between photoperiods, i4E samples formed a single cluster. In i4L, a greater transcriptomic divergence was observed, but this was lost again upon molting into i5E, the samples of which clustered very closely with i4E libraries. This pattern suggests that the transcriptional divergence between 12L:12D and 18L:6D individuals at i4L, which presumably generates the differences in size between instar five wing buds (Figure 1D), are effectively reset upon the moult to instar five. Finally, i5L showed a much clearer and more extensive divergence with respect to all previous developmental stages, as well as between photoperiods. This pattern suggests that the regulatory mechanism(s) of wing length determination as a response to day length are activated sometime between i5E and i5L.

**Figure 2.**
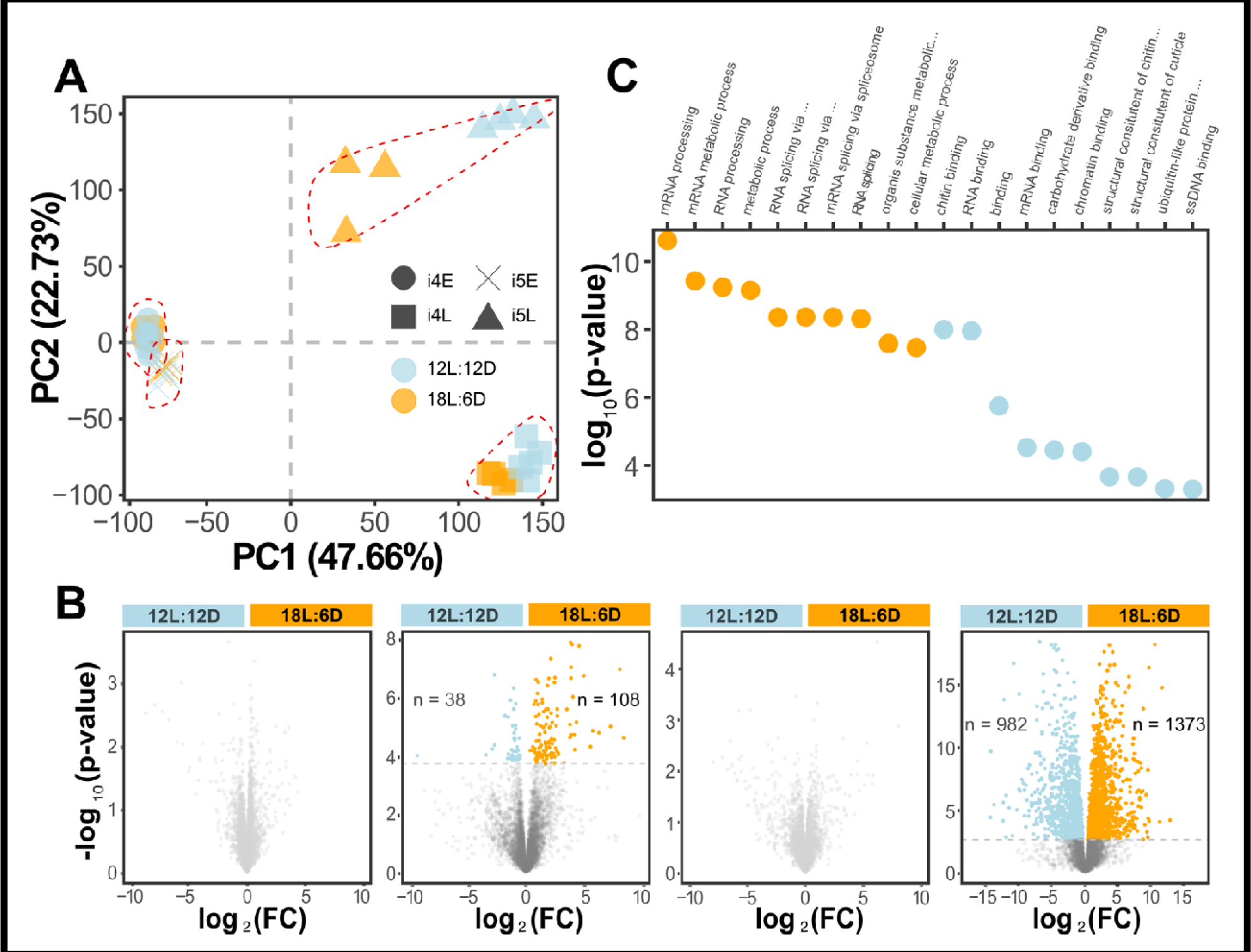
Extensive differential expression induced by photoperiod throughout development. **A**) PCA plot resulting from wing transcriptomic data obtained from four developmental timepoints (i4E, i4L, i5E, i5L) between photoperiods. **B)** Volcano plots for in between-photoperiod comparisons for, left to right, i4E, i4L, i5E, i5L. Grey dots represent non-differentially expressed genes. Blue dots represent gene with significantly higher expression in 12L:12D. Orange dots represent genes with significantly higher expression in 18L:6D. Horizontal grey lines indicate the threshold p-value equivalent to FDR = 0.01 after correction. **C)** Most enriched Biological Process (orange) and Molecular Function (blue) GO Terms in i5L based on the differentially expressed genes.

### Extensive differences in wing bud gene expression in response to photoperiod

Differential expression analysis was consistent with the trends observed in the PCA plots (Figure 2B). The low-expression filtering resulted in the exclusion of about two thirds of the total annotated genes (n = 20,431), and 7,361 genes being retained for DE analysis. In i4E and i5E, no genes were differentially expressed between 12L:12D and 18L:6D (Figure 2B), in agreement with the tight clustering of samples perceived in the PCA (Figure 2A). As opposed to these early stages into nymphal instars, 146 genes were differentially expressed between photoperiods in i4L, of which 38 were upregulated in 12L:12D and 108 in 18L:6D (Figure 2B, Supplemental Data File 1). At i5L, in agreement with the more divergent response to photoperiod at that stage perceived in the PCA (Figure 2A), 2355 genes were differentially expressed; 982 were upregulated in 12L:12D and 1373 in 18L:6D (Figure 2B, Supplemental Data File 2).

These differentially expressed genes at both i4L and i5L were used to run GO Term enrichment tests, in order to explore which processes responded differently to the alternative photoperiod treatments, with the ultimate goal of identifying candidate molecular pathway responsible for the regulation of wing formation and development. In i4L, the analysis revealed a total of 344 significantly enriched biological process (BP) GO Terms, with the 10 most enriched terms mostly associated with cell cycle regulation, DNA replication and RNA transcription (Figure S4A). Interestingly, no terms associated with specific wing or imaginal disc growth were among them (Supplemental Data File 3), suggesting that the differential expression triggered by photoperiod differences are involved in systemic regulation of cell division but without translating into tissue-specific wing development and patterning. Simultaneously, there were 66 molecular function (MF) GO Terms which showed a significant enrichment in i4L, mostly associated with helicase activity (Figure S4B). The bigger DE dataset in i5L, on the other hand, generated 729 significantly enriched BP and 156 MF GO Terms. While the 10 most-enriched BP terms were mostly associated with RNA processing and splicing (Figure 2C), the list also included several terms associated with wing and imaginal disc development (Figure S5A). These results highlight that differential wing growth is at least in part mediated through differential expression of genes known to act in insect wing development. Interestingly, several of the most enriched MF terms in i5L were associated with DNA and RNA binding, in concordance with the BP top-hits and suggestive of differences in the modulation of transcription factors and splicing regulators (Figure 2C).

We further explored the significantly enriched ontology to narrow down the list of candidate pathways responsible for the regulation of wing growth in *G. buenoi*, based on previous findings in other insect species (Gotoh *et al*., 2015; Hayes *et al*., 2019; Zhang, Brisson and Xu, 2019; Tripathi and Irvine, 2022). Intriguingly, we found a significant enrichment for genes involved in both ecdysone signaling and the Hippo pathway (Figure S5, Supplemental Data File 4), as well as insulin, highlighting its importance in growth regulation despite playing no specific role in wing polyphenism in *G. buenoi* (Gudmunds *et al*., 2022). In total, 28 out of 105 involved in the Hippo signaling pathway were differentially expressed in i5L. In particular, the genes Fat (Ft; logFC = 0.92, FDR = 5.97e-5) and Dachsous (Ds; logFC = 2.07, FDR = 1.69e-7) i.e., core components of one of the regulatory branches of Hippo signaling (Gridnev and Misra, 2022), as well as the transcriptional co-activator Yorkie (Yki; logFC = 3.26, FDR = 1.68e-4), were upregulated in 12L:12D (Figure 3A, Supplemental Data File 2). In 18L:6D, multiple genes were upregulated too, including Dachs (D; logFC = 1.01, FDR = 6.06e-3), which links Fat/Ds activity to the core Hippo kinase cassette (Mao *et al*., 2006), and 14-3-3 (logFC = 1.40, FDR = 1.52e-5), the protein responsible for sequestering phosphorylated Yki in the cytosol (Misra and Irvine, 2018).

**Figure 3.**
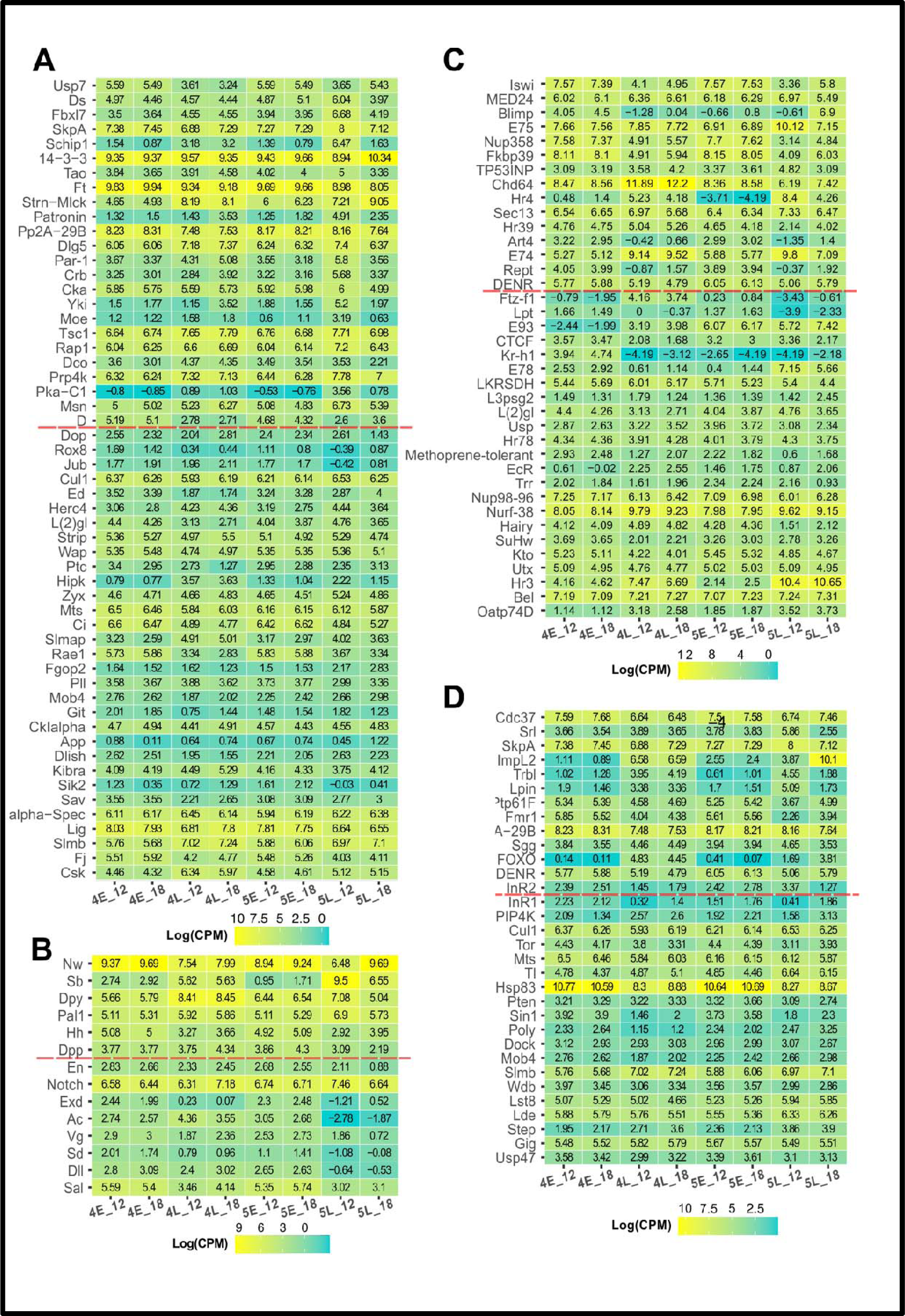
Normalized expression dynamics of genes belonging to or modulated by the Hippo signaling pathway (A), involved in wing formation and development (B), belonging to or modulated by ecdysone signaling (C), and belonging to or modulated by insulin (D). Genes are ordered top to bottom by FDR value in the comparison between photoperiods at i5L. Genes located above the red dotted line present significantly different expression levels. All indicated values are log_2_ of the average CPM for each given treatment.

To better understand the expression dynamics of Ft, Ds and Yki over the course of wing development, we examined CPM values for all sampled time points in both photoperiods (Figure 3A). In i4L, Yki showed differential expression between photoperiods, with a higher expression in 18L:6D (logFC = 2.40, FDR = 6.50e-3; Figure 3A). Later in i5L, the expression pattern was reversed, with an increase in expression specific to individuals reared in 12L:12D, while remaining unchanged in 18L:6D. This is likely the reflection of a high degree of cell proliferation in 12L:12D, an indispensable requisite for the generation of a macropterous wing. Ft expression was highest in the early stages within each instar. With respect to i5E, the expression levels in i5L decreased in both photoperiods, but the magnitude of decrease was lower in 12L:12D, generating distinct differences in Ft expression at i5L between the two photoperiods (Figure 3A). Finally, Ds levels remained relatively stable in both photoperiods across development until i5L, where the level decreased in 18L:6D but increased in 12L:12D with respect to i5E (Figure 3A). These in-depth exploration of the expression levels of the genes belonging to Fat/Hippo signaling evidences that the core genes of the pathway exhibit dynamic patterns of expression in nascent wings of *G. buenoi*. However, these expression patterns are gene-specific and variable, even among the genes of the same signaling pathway. In addition to the genes highlighted above, we did not find any of the core Hippo pathway kinases e.g., Hippo, Warts, Salvador or Mats, to be differentially expressed in i5L (Supplemental Data File 2).

Among the differentially expressed genes at i5L, we also identified three core genes involved in wing size regulation in *Drosophila* (Figure 3B). In particular, Dumpy (Dpy; logFC = 2.04, FDR = 5.07e-6) and Peptidyl-α-hydroxyglycine-α-amidating lyase 1 (Pal1; logFC = 1.17, FDR = 6.46e-6) generate smaller or shorter wings when knocked-down with RNAi, whereas Narrow (Nw; logFC = 3.20, FDR = 1.71e-10) RNAi generates a longer narrower wing (Ray *et al*., 2015). The upregulation of Dpy and Pal1 in 12L:12D and Nw in 18L:6D suggests that the activity of these genes is shared between *G. buenoi* and *D. melanogaster*.

### Gene coexpression network across development between photoperiods

The expression dynamics of DE genes involved in the Hippo signaling pathway and wing size regulation was extremely labile i.e., the absolute expression level and in which treatment (developmental stage and photoperiod) this was observed varied substantially (Figure 3A, B). Additionally, many genes that did not show evidence of DE might still play important roles in wing tissue development through more subtle differences in transcript abundance depending on the environmental conditions. These two factors motivated the necessity of exploring genome-wide patterns of expression using WGCNA. The gene correlation network analysis identified 13 co-expression clusters among the 7,361 total genes that were retained after the low-expression filtering step. Out of those, 5,519 genes were included in one of the clusters (Figure 4A, Table S4). We observed, overall, that developmental progression is the main driver of differences in co-expression dynamics i.e., most of the changes in eigenvalue correlation happen as a consequence of differences in developmental time, while relative differences in response to photoperiod are more restricted to specific modules (Figure 4A, S6). In particular, co-expression modules 2 and 5, and to a lesser extent modules 1 and 4, showed considerable differences in eigenvalue correlation between both photoperiods in i5L (Figure 4A, S6).

All four modules showed radically different gene trajectories across development, although they all shared a general similarity between photoperiod treatments in the earliest three time points (i.e., before reaching i5L). Module 1 was characterized by high relative expression levels in i4E and i5E, with reductions in transcript abundance in i4L and more apparent differences in i5 (Figure 4B). Modules 2 and 5 shared low relative expression levels of their genes until i5L, when genes belonging to module 2 showed a significantly higher expression in 12L:12D, while those included in module 5 were more highly expressed in 18L:6D. (Figure 4C, E). On the other end, module 4 did not show drastic differences in transcript abundance except in i4L, when 12L:12D had on average higher expression (Figure 4D).

**Figure 4.**
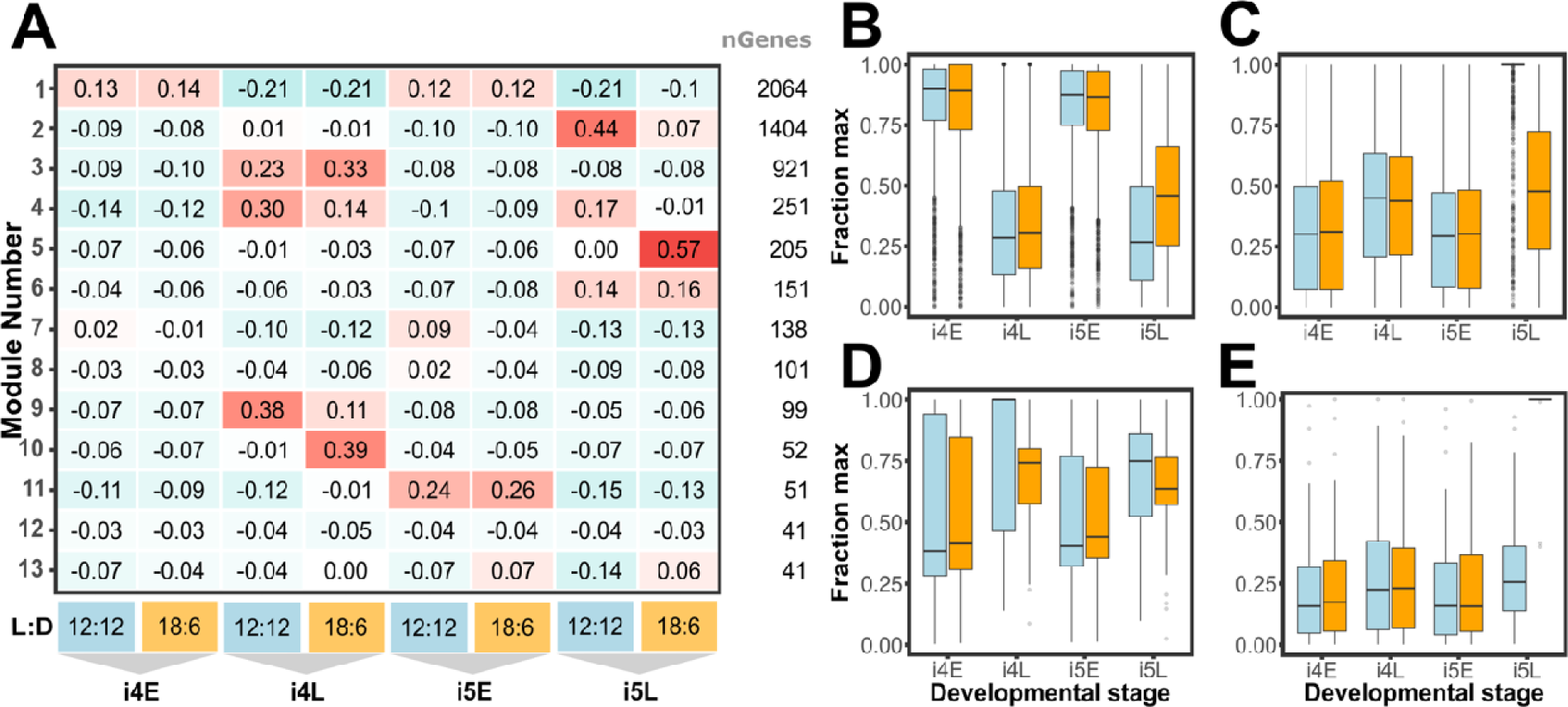
WGCNA of all differentially expressed genes across development and between photoperiods. **A**) Correlation of the eigengene for each cluster with the eight treatments (four developmental stages, 2 photoperiods). Modules are ordered by number of genes they include. Th indicated values represent the median across all libraries corresponding to the same developmental stage and photoperiod. **B-E)** Expression trajectories of the genes clusters that showed the highest difference between photoperiods in i5, including module 1 **(B)**, 2 **(C)**, 4 **(D)** and 5 **(E)**. For each gene, the average expression per treatment was calculated, and all expression levels are shown relative to this maximum. Blue columns indicate the gene expression patterns at 12L:12D, and orange at 18L:6D.

We functionally assessed these modules with *topGO* and found a significant enrichment for genes involved in Hippo signaling (p = 1.9e-3) in module 2 (Supplemental Data File 6), while module 1 was enriched in genes positively regulating ecdysone signaling processes (p = 0.026, Supplemental Data File 5). Regarding the involvement of the different co-expression modules with wing formation, module 2 showed a significant enrichment for genes associated with wing disc morphogenesis (GO:0007472; p = 0.031) and development (GO:003522; p = 0.006). Finally, module 1 was heavily enriched in genes associated with mRNA splicing and its regulation (Supplemental Data File 5).

### RNAi on candidate genes for wing length determination

From the list of genes that were differentially expressed in i5L (Supplemental Data File 2) we selected a subset of candidate genes for functional validation which we based on the observed expression patterns and also earlier annotation pointing to wing development in *Drosophila* or involvement in hormonal regulation (see Supplemental Table S5 for specific motivations for each gene). In the screen we injected dsRNA during late stages of instar four in order for the RNAi effect to be initiated before or as early as possible during instar five (see methods for further details). None of the instar four RNAi treatments against the genes listed in Supplemental Table S5 had a clear effect on adult wing morph frequencies (data not shown), suggesting that these genes either have no role in wing morph determination or that the knockdown failed or was not sufficiently strong to provoke a response during adult wing development. Interestingly, individuals treated with Ds or Ft dsRNA generated two similar but distinct wing bud shape phenotypes after the moult to instar five (Figure 5A). Ft RNAi wing buds had a pointier tip compared to dsGFP individuals, and Ds RNAi generated wing buds with a blunter and shorter appearance than normal wing buds (Figure 5A). Double knockdown of Ft and Ds produced wing buds with an intermediate phenotype, appearing blunter than Ft wing buds but shorter and less wide than Ds wing buds (Figure 5A). These results show that Ft and Ds act to shape the instar five wing bud which forms during the fourth instar. The adult wings of Ft, Ds, or Ft/Ds RNAi individuals injected in instar four were generally as long as dsGFP wings but were broader.

**Figure 5.**
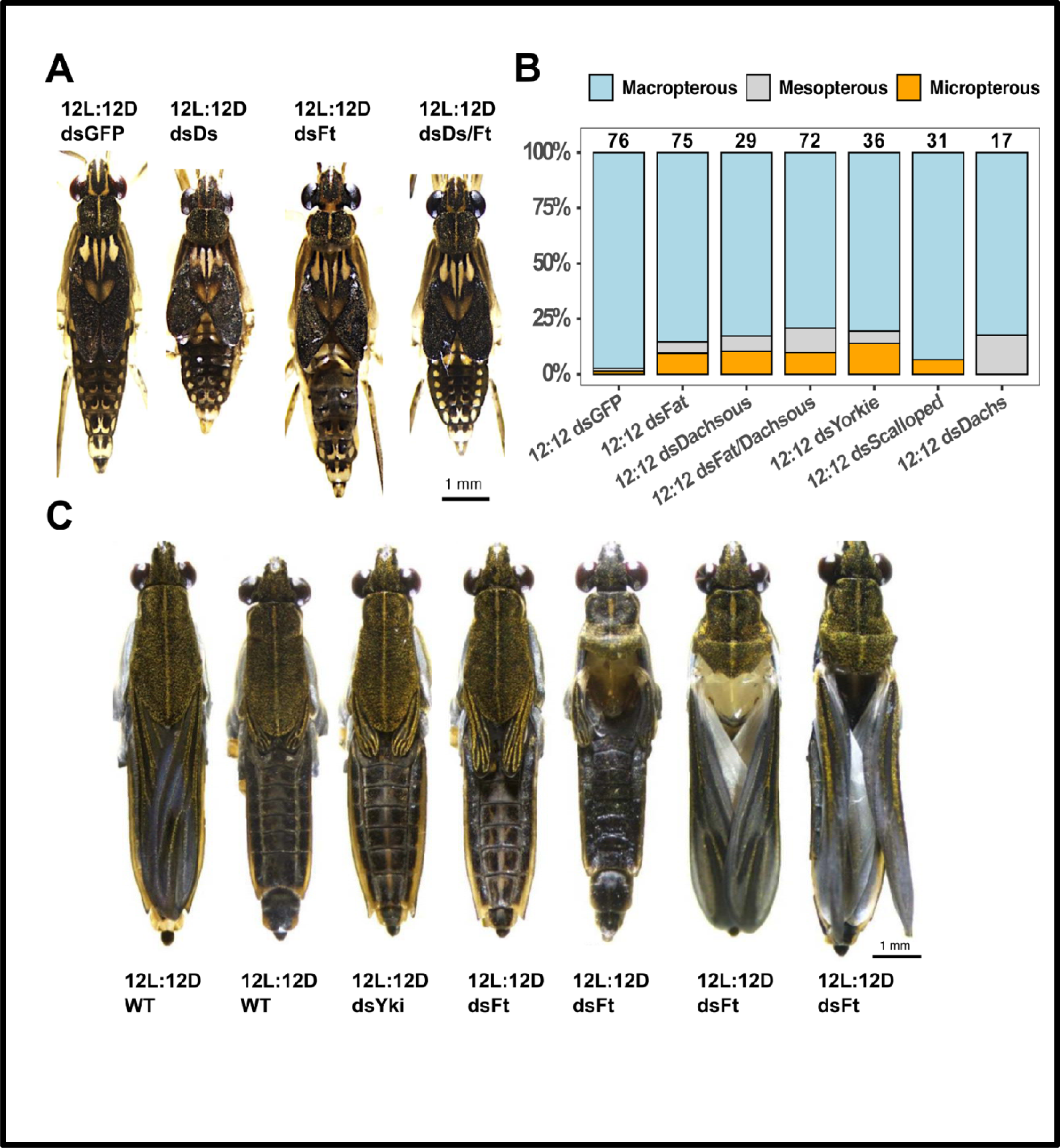
RNAi against genes in the Fat/Hippo pathway causes wing bud phenotypes and wing morph switches. **A**) Phenotypes of i5 individuals treated with Ds, Ft or Ds/Ft RNAi during i4. **B)** Adult wing morph frequencies of individuals treated with RNAi against genes in the Fat/Hippo pathway. **C)** Adult individuals, either wild type reared in 12L:12D or 18L:6D, or treated with dsYki or dsFt in 12L:12D. The displayed dsYki and dsFt RNAi phenotypes are representative of the range of phenotype observed after RNAi against Yki, Ds, Ft, Sd and D.

### Fat/Dachsous/Yki differential expression causes wing length variation in G. buenoi

Since the Fat/Hippo pathway seem to control wing size in both Holometabolous (Gridnev and Misra, 2022) and Hemimetabolous (Ohde, Mito and Niimi, 2022) insects and that we found a role of the Fat/Hippo signaling components Ds and Ft in wing bud development as manifested by the knockdowns initiated in instar four, we decided to explore their roles in *G. buenoi* wing development further. To do so we injected Ds and Ft early in instar five as opposed to the instar four injections performed in the screen of candidate genes. Here, RNAi against Ds and Ft resulted in a significantly higher proportion of micropterous and mesopterous individuals compared to the dsGFP control (Ds: χ^2^ = 5.0, d.f. = 1, *P* = 0.02, Ft: χ^2^ = 5.5, d.f. = 1, *P* = 0.02, Figure 5B). The same outcome was obtained when simultaneously knocking down Ds and Ft (χ^2^ = 10.3, d.f. = 1, *P* = 0.001). Furthermore, whereas high doses of Yki dsRNA were lethal when injected during instar four, lower doses given to early in instar five led to less mortality and thus we were able to score wing morphs in the adults. When injecting individuals in 12L:12D photoperiod we found a significantly lower frequency of long-winged individuals (χ^2^ = 7.2, d.f. = 1, *P* = 0.007; Figure 5B), in accordance with the central role for Yki in the Fat/Hippo signaling pathway. Yki is a transcriptional co-regulator and thus does not bind to DNA itself, rather it interacts with the transcription factor Scalloped (Sd). We thus investigated the role of Sd in wing morph determination, but RNAi against Sd did not have a statistically significant effect on wing morph frequencies (χ^2^ = 0.15, d.f. = 1, *P* = 0.70; Figure 5B). We also targeted Dachs (D) with RNAi, the role of this protein is to mediate the signaling between Ft/Ds to the core kinase cassette of the Hippo pathway (Mao *et al*., 2006). Here we knocked down D expression in both 18L:6D and 12L:12D, since D was found to be significantly higher expressed in 18L:6D. RNAi against this gene generated an elevated frequency of the mesopterous morph in 12L:12D whereas it had no clear effect in 18L:6D. The wing morph frequencies between dsGFP and dsDachs treated individuals in 12L:12D was near statistical significance (χ^2^ = 3.6, d.f. = 1, *P* = 0.06; Figure 5B). For the genes producing significant changes in wing morph frequencies, we observed significant reduction in mRNA levels after RNAi treatments using RT-qPCR (Figure S7).

The wings of RNAi-affected individuals resembled to a large degree normal micropterous or mesopterous wings (Figure 5C). However, all genes that showed an effect on wing morph frequencies also had an effect on the pronotum, which were short and wrinkled. This phenotype appeared to interfere with proper positioning of the wings along the dorsal abdomen (Figure 5C) but was not present in all individuals that switched wing morphs due to the RNAi treatment.

## Discussion

Wing polyphenism in insects has for long been a subject of interest from an evolutionary and ecological perspective (Järvinen and Vepsäläinen, 1976; Vepsäläinen, 1978; Harrison, 1980; Roff, 1986; Fairbairn, 1988; Spence, 1989; Andersen, 1993; Harada and Numata, 1993; Ahlroth *et al*., 1999) and this interest has only increased in recent years with the new possibilities that molecular genetics confers (Hayes *et al*., 2019; Zhang, Brisson and Xu, 2019). Indeed, a few studies have now been able to identify the regulatory network involved in nutritionally-induced wing polyphenism as exemplified by the IIS pathway in some hemipterans (Xu *et al*., 2015; Fawcett *et al*., 2018; Smýkal *et al*., 2020). It is clear however that this is not a common route in all hemipterans to the evolution of wing polyphenism (Gudmunds *et al*., 2022). In the present study we aimed to identify the gene regulatory network which control photoperiod induced wing morph determination in the water strider *G. buenoi* by examining wing transcriptomic differences underlying development of wing morphs in direct association to their respective inductive environments. Our results demonstrate that the conserved Fat/Hippo pathway controls wing length polyphenism in *G. buenoi* and provides an important basis for future comparative studies that examine how different environmental cues can be sensed and co-opted with different cellular and neuroendocrine pathways to control environmentally induced phenotypes.

### Transcriptomic approaches to study wing length polyphenism

The ability to grow long or short wings in response to environmental cues is a prevalent trait in insects (Roff, 1986; Zera and Denno, 1997; Hayes *et al*., 2019; Zhang, Brisson and Xu, 2019). Our understanding of the molecular basis of this trait was for decades limited, but has improved in the recent years due to the development of computational approaches such as RNA-seq analysis (Xu and Zhang, 2017; McCulloch *et al*., 2019) and functional studies using RNAi (Zhang, Brisson and Xu, 2019). The majority of studies that have functionally verified the role of specific genes in wing length determination have selected and tested candidate genes based on *a priori* functional knowledge from literature on a gene’s importance in growth regulation and connection to environmental factors. This approach has proven fruitful as manifested in the proliferation of studies showing that the IIS pathway is regulating wing morph determination in at least three Hemipterans to date (Xu *et al*., 2015; Fawcett *et al*., 2018; Smýkal *et al*., 2020). Nonetheless, unbiased approaches to identify candidate genes have been lacking, likely due to a difficulty to predict adult wing morph due to incomplete penetrance of the response to environmental cues.

In the brown planthopper, two recent studies using morph-specific RNAseq have nevertheless gained new interesting insights on the genetic mechanism of wing morph induction (Zhang *et al*., 2021, 2022). For example, the studies revealed that FOXO, being central for wing morph determination, regulates Vg expression by binding to introns within the *vg* locus and that the Zfh1 transcription factor regulates FOXO expression (Zhang *et al*., 2021, 2022). However, the prediction of wing morph in these studies was achieved by using RNAi, and thus lacks a clear connection to the environmental conditions controlling wing morph determination. Therefore, it becomes difficult to control for artefactual gene expression due to the RNAi treatment as well as to connect the local response in wing tissue to the action of systemic hormone signaling. The robust predictability of *G. buenoi* wing morph based on photoperiod during nymphal development (Gudmunds *et al*., 2022) overcomes these hurdles. It allows for the combined use of RNA-seq analysis to explore the effects of the inductive environmental cue at a transcriptome-wide level, and in turn facilitates an unbiased determination of candidate genes for further verification with RNAi. Overall, it opens the door to explore the proximate genetic mechanisms of wing polyphenism without direct manipulation of gene expression, and directly connects the findings to the environmental cue.

### Gene expression divergence driven by photoperiod varies over wing development progression

In the present study, we observe dynamic gene expression profiles, both throughout development and between photoperiods (Figure 2A, B).

In i4L, 146 genes were differentially expressed, although they do not seem to be directly involved in wing length determination, judging by the lack of ontology enrichment in genes associated with imaginal disc and wing development (Figure S4). These differentially expressed genes, however, might be relevant for the determination of wing bud size upon moulting to i5. In i5L, the effect of differences in photoperiod becomes more apparent, with almost a third of all expressed genes show differential levels of transcript abundance between 12L:12D and 18L:6D. In contrast, at i4E and i5E we find no differentially expressed genes (Figure 2B). The lack of differentially expressed genes at i5E is particularly interesting given that some transcriptional divergence was detected at i4L. These patterns of gene expression between wing morphs are similar to those found in the brown planthopper (Zhang *et al*., 2021). In particular, Zhang and colleagues found that only one single gene was differentially expressed between wing morphs at 24 hae, corresponding to i5E in our experiment (Zhang *et al*., 2021). Together, these data suggest that a general pattern of wing polyphenic development in hemipterans is that at the onset of development to the alternative wing morphs there are no detectable traces of the transcriptional cascades leading up to morph differentiation. It is thus likely that the effect of hormonal regulators is initiated after 24 hae and drive wing morph development in a relatively short time-window before moulting to adulthood.

The hormonal control of wing morph determination is not yet known in *G. buenoi*. In the brown planthopper both JH and insulin-like peptides are implicated in wing morph determination (Xu *et al*., 2015; Ye *et al*., 2019). In *G. buenoi*, despite the differential expression of multiple insulin-associated genes (Figure 3D), we recently showed that insulin signaling is not involved in the regulation of wing morphology. Here, where photoperiod is the main environmental cue for induction of wing morphs, we find 20E to be a likely candidate endocrine regulator since its release/synthesis can be regulated by photoperiod (Steel and Vafopoulou, 2006) and has been linked to photoperiod-controlled polyphenisms before (Nijhout, 2009). This idea is supported by the differentially expressed genes in i5L, where gene ontology analysis revealed an enrichment for the regulation of ecdysone-receptor mediated signaling (Figure S5). Among the genes causing this enrichment we find E75, E74 and Hr4 (Figure 3C) – all well-known ecdysone-responsive genes that act as transcription factors to mediate the large transcription cascades that are induced in tissues and cell in the presence of ecdysone (Uyehara and McKay, 2019; Uyehara, Leatham-Jensen and McKay, 2022).

We hypothesize that the differential expression of these genes is a signature of different ecdysone titers in the alternative wing morphs, which in turn are caused by the exposure to the different photoperiods. While this hypothesis needs further exploration, it is likely that variable ecdysone titers over development are inducing the gene expression programs in the growing wing tissue that lead to development of the two wing morphs, similar to polyphenic mechanisms acting in other insects (Rountree and Nijhout, 1995; Lobbia, Niitsu and Fujiwara, 2003; Bhardwaj *et al*., 2020; van der Burg *et al*., 2020). Together with the regulation of wing length, the variable ecdysone titers and all the responsive genes are likely responsible for the regulation of many correlated traits (Roff, 1984; Zera, 1984; Kaitala and Huldén, 1990; Crnokrak and Roff, 1995; Zera and Larsen, 2001) in other tissues, although we do not capture these associated variations in our tissue-specific transcriptome approach.

### The Fat/Hippo signaling pathway mediates wing polyphenism G. buenoi

The Fat/Hippo signaling pathway plays an important role in growth regulation of *Drosophila* wings (Irvine and Harvey, 2015; Gou, Lin and Othmer, 2018; Gridnev and Misra, 2022) and RNAi phenotypes of Ft (Hust *et al*., 2018; Ohde, Mito and Niimi, 2022) or Ds (Gotoh *et al*., 2015) generates individuals with disproportionally shorter wings in both Hemimetabolous and Holometabolous insects, suggesting a conserved role in regulation of wing size. Furthermore, the Fat/Hippo pathway has been proposed to be a missing link in connecting circulating signals (i.e. hormones) to localized tissue responses (Gotoh *et al*., 2015). In *Drosophila*, ecdysone signaling and the Fat/Hippo pathway are functionally connected through the ability of EcR and Yki to interact via the EcR co-activator Taiman (Zhang *et al*., 2015). It was therefore intriguing to find multiple members of this pathway to be differentially expressed in nascent wing tissue, with a significant enrichment of Hippo signaling and its regulation in the gene ontology analysis (Figure S5). Notably, the core components Ft, Ds and Yki all showed significantly higher expression in 12L:12D than in 18L:6D at the latest i5L timepoint. RNAi against Ft and Ds in instar four generated individuals with abnormal wing bud phenotypes, showing that Ft and Ds signaling is important during instar four to correctly shape wing progenitor structures (Figure 5C). In contrast, RNAi against Ft, Ds and Yki initiated in instar five generated no distinct abnormalities in adult wing shape, instead, a proportion of affected individuals appeared with short wings, phenocopying the mode of wing development occurring in 18L:6D. The penetrance of this phenotype was only ∼15-25% but nonetheless statistically significant for Ft, Ds, the cocktail Ft/Ds and Yki, thus strongly indicating a causative role for Fat/Hippo signaling in *G. buenoi* wing polyphenism. Although we focused on these three genes to functionally verify their role in regulating the polyphenism, it is noteworthy that genes in the Fat/Hippo pathway did not all follow the same expression dynamics across development and as a response to photoperiod. This is evidenced by the results of WGCNA, as four different gene modules showed considerable differences in i5L (Figure 4A). Module 2 was the only one enriched for the Fat/Hippo pathway, and was characterized by low expression levels through development and a sharp increase in i5L, accentuated in 12L:12D (Figure 4C). This however was not the case for all genes, and some showed opposite trajectories throughout development, including Ft (Figure 4A).

The way by which Fat/Hippo signaling regulates development and growth is relatively well-known through research in *Drosophila* wing discs (see Gridnev and Misra, 2022 for a recent review). Here, Ft is uniformly expressed in the wing disc whereas Ds is expressed in a gradient and the relative steepness of this gradient, together with that of other Fat/Hippo signaling components, is in control of Yki activity and growth (Matakatsu and Blair, 2006; Rogulja, Rauskolb and Irvine, 2008; Willecke *et al*., 2008). The establishment of the Ds gradient occurs through the influence of Vg (Gridnev and Misra, 2022). It is noteworthy that the core genes regulating wing patterning in *Drosophila* (Notch, wingless, engrailed, vestigial and cut; Neumann and Cohen, 1996; Tripathi and Irvine, 2022) do not show evidence of DE in i5L in *G. buenoi* (Figure 4B), but the morphogen decapentaplegic (Dpp), which presents a gradient in abundance that orchestrates wing growth through the Fat/Hippo signaling cascade (Rogulja, Rauskolb and Irvine, 2008; Bosch *et al*., 2017; Tripathi and Irvine, 2022) together with its regulator hedgehog (hh; Tanimoto *et al*., 2000) are differentially expressed. These results are in line with the observation that long and short wings are structurally equivalent structures which only vary in absolute size. We hypothesize that the observed differential regulation of Ft and Ds between photoperiods could provide the means to regulate wing size by establishing different gradients of Ft/Ds expression. One that sustains high levels of cell proliferation, aided by a high expression of Yki, forming macropterous wings (12L:12D) and another that restricts proliferation (18L:6D). Additionally, the lower expression and activation of Yki in nascent micropterous wings may result in de-repression of genes involved in programmed cell death (Verghese, Bedi and Kango-Singh, 2012) which could play a role in the formation of a micropterous wing. How regulation of expression levels of Ds, Ft and Yki is achieved and whether gradient expression of Ds occurs in *G. buenoi* wings is an interesting avenue for further research. Given that the differentially expressed genes in i5L shows an enrichment for genes involved in ecdysone signaling, it is likely that ecdysone is acting on the wing tissue at this timepoint to program development and growth, and is thus a highly interesting candidate hormone to generate the differential regulation of genes in the Fat/Hippo pathway.

## Conclusions

Here we used RNAseq to identify the gene regulatory network involved in the control of wing polyphenism in the water strider *G. buenoi*. Interestingly, several genes in the conserved Fat/Hippo signaling pathway were differentially expressed, including Ft, Ds and Yki. When these genes were silenced with RNAi in individuals destined to become macropterous, a significant proportion of individuals instead appeared with micropterous or mesopterous wings. Therefore, we conclude that the Fat/Hippo pathway is involved in the regulation of wing polyphenism in *G. buenoi* in response to different photoperiods. Future research will be needed to understand how Fat/Hippo signaling is responding to differences in photoperiod in *G. buenoi*.

## Acknowledgements

The authors acknowledge support from the National Genomics Infrastructure in Stockholm funded by Science for Life Laboratory, the Knut and Alice Wallenberg Foundation and the Swedish Research Council, and SNIC/Uppsala Multidisciplinary Center for Advanced Computational Science for assistance with massively parallel sequencing and access to the UPPMAX computational infrastructure.

## References

1. Ahlroth, P. et al. (1999) ‘Geographical variation in wing polymorphism of the waterstrider Aquarius najas (Heteroptera, Gerridae)’, Journal of Evolutionary Biology, 12, pp. 156–160.

2. Alexa, A. and Rahnenfuhrer, J. (2023) ‘topGO: Enrichment Analysis for Gene Ontology’. Bioconductor version: Release (3.16). Available at: 10.18129/B9.bioc.topGO.

3. Andersen, N.M. (1982) The Semiaquatic Bugs (Hemiptera, Gerromorpha) – Phylogeny, Adaptations, Biogeography and Classification. Klampenborg, Denmark: Scandinavian Science Press LTD. (Entomonograph, Volume 3).

4. Andersen, N.M. (1993) ‘The Evolution of Wing Polymorphism in Water Striders (Gerridae): A Phylogenetic Approach’, Oikos, 67(3), pp. 433–443. Available at: 10.2307/3545355.

5. Armisén, D. et al. (2018) ‘The genome of the water strider Gerris buenoi reveals expansions of gene repertoires associated with adaptations to life on the water’, BMC Genomics, 19(1), p. 832. Available at: 10.1186/s12864-018-5163-2.

6. Atkinson, D. (1994) ‘Temperature and Organism Size—A Biological Law for Ectotherms?’, in M. Begon and A.H. Fitter (eds) Advances in Ecological Research. Academic Press, pp. 1–58. Available at: 10.1016/S0065-2504(08)60212-3.

7. Bakker, K. (1959) ‘Feeding Period, Growth, and Pupation in Larvae Ofdrosophila Melanogaster’, Entomologia Experimentalis et Applicata, 2(3), pp. 171–186. Available at: 10.1111/j.1570-7458.1959.tb00432.x.

8. Bhardwaj, S., et al. (2020) ‘Origin of the mechanism of phenotypic plasticity in satyrid butterfly eyespots’, eLife. Edited by P.J. Wittkopp, K.E. Sears, and C. Wheat, 9, p. e49544. Available at: 10.7554/eLife.49544.

9. Bosch, P.S. et al. (2017) ‘Dpp controls growth and patterning in Drosophila wing precursors through distinct modes of action’, eLife, 6, p. e22546. Available at: 10.7554/eLife.22546.

10. Brogiolo, W. et al. (2001) ‘An evolutionarily conserved function of the Drosophila insulin receptor and insulin-like peptides in growth control’, Current Biology, 11(4), pp. 213–221. Available at: 10.1016/S0960-9822(01)00068-9.

11. Brůna, T. et al. (2021) ‘BRAKER2: automatic eukaryotic genome annotation with GeneMark-EP+ and AUGUSTUS supported by a protein database’, NAR Genomics and Bioinformatics, 3(1), p. lqaa108. Available at: 10.1093/nargab/lqaa108.

12. van der Burg, K.R.L. et al. (2020) ‘Genomic architecture of a genetically assimilated seasonal color pattern’, Science, 370(6517), pp. 721–725. Available at: 10.1126/science.aaz3017.

13. Camacho, C. et al. (2009) ‘BLAST+: architecture and applications’, BMC Bioinformatics, 10(1), p. 421. Available at: 10.1186/1471-2105-10-421.

14. Crnokrak, P. and Roff, D.A. (1995) ‘Fitness differences associated with calling behaviour in the two wing morphs of male sand crickets, *Gryllus firmus*’, Animal Behaviour, 50(6), pp. 1475– 1481. Available at: 10.1016/0003-3472(95)80004-2.

15. Danecek, P. et al. (2021) ‘Twelve years of SAMtools and BCFtools’, GigaScience, 10(2), p. giab008. Available at: 10.1093/gigascience/giab008.

16. Dobin, A. et al. (2013) ‘STAR: ultrafast universal RNA-seq aligner’, Bioinformatics, 29(1), pp. 15–21. Available at: 10.1093/bioinformatics/bts635.

17. Fairbairn, D.J. (1988) ‘Adaptive significance of wing dimorphism in the abscence of dispersal: a comparative study of wing morphs in the waterstrider, *Gerris remigis*’, Ecological Entomology, 13, pp. 273–281.

18. Fawcett, M.M. et al. (2018) ‘Manipulation of insulin signaling phenocopies evolution of a host-associated polyphenism’, Nature Communications, 9(1), p. 1699. Available at: 10.1038/s41467-018-04102-1.

19. Fernandez-Nicolas, A. et al. (2022) ‘Broad complex and wing development in cockroaches’, Insect Biochemistry and Molecular Biology, 147, p. 103798. Available at: 10.1016/j.ibmb.2022.103798.

20. Flynn, J.M. et al. (2020) ‘RepeatModeler2 for automated genomic discovery of transposable element families’, Proceedings of the National Academy of Sciences, 117(17), pp. 9451–9457. Available at: 10.1073/pnas.1921046117.

21. Gabriel, L. et al. (2021) ‘TSEBRA: transcript selector for BRAKER’, BMC Bioinformatics, 22(1), p. 566. Available at: 10.1186/s12859-021-04482-0.

22. Gokhale, R.H. et al. (2016) ‘Intra-organ growth coordination in Drosophila is mediated by systemic ecdysone signaling’, Developmental Biology, 418(1), pp. 135–145. Available at: 10.1016/j.ydbio.2016.07.016.

23. Gotoh, H. et al. (2015) ‘The Fat/Hippo signaling pathway links within-disc morphogen patterning to whole-animal signals during phenotypically plastic growth in insects’, Developmental Dynamics, 244(9), pp. 1039–1045. Available at: 10.1002/dvdy.24296.

24. Gou, J., Lin, L. and Othmer, H.G. (2018) ‘A Model for the Hippo Pathway in the Drosophila Wing Disc’, Biophysical Journal, 115(4), pp. 737–747. Available at: 10.1016/j.bpj.2018.07.002.

25. Gramates, L.S. et al. (2022) ‘Fly Base: a guided tour of highlighted features’, Genetics, 220(4), p. iyac035. Available at: 10.1093/genetics/iyac035.

26. Gridnev, A. and Misra, J.R. (2022) ‘Emerging Mechanisms of Growth and Patterning Regulation by Dachsous and Fat Protocadherins’, Frontiers in Cell and Developmental Biology, 10, p. 842593. Available at: 10.3389/fcell.2022.842593.

27. Gudmunds, E. et al. (2022) ‘Photoperiod controls wing polyphenism in a water strider independently of insulin receptor signalling’, Proceedings of the Royal Society B: Biological Sciences, 289(1973), p. 20212764. Available at: 10.1098/rspb.2021.2764.

28. Harada, T. and Numata, H. (1993) ‘Two critical day lengths for the determination of wing forms and the induction of adult diapause in the water strider, *Aquarius paludum*’, Naturwissenschaften, 80, pp. 430–432.

29. Harrison, R.G. (1980) ‘Dispersal Polymorphisms in Insects’, Annual Review of Ecology and Systematics, 11(1), pp. 95–118. Available at: 10.1146/annurev.es.11.110180.000523.

30. Hayes, A.M., et al. (2019) ‘Mechanisms regulating phenotypic plasticity in wing polyphenic insects’, in R. Jurenka (ed.) Advances in Insect Physiology. Academic Press, pp. 43–72. Available at: 10.1016/bs.aiip.2019.01.005.

31. Herboso, L. et al. (2015) ‘Ecdysone promotes growth of imaginal discs through the regulation of Thor in D. melanogaster’, Scientific Reports, 5(1), p. 12383. Available at: 10.1038/srep12383.

32. Hietakangas, V. and Cohen, S.M. (2009) ‘Regulation of Tissue Growth through Nutrient Sensing’, Annual Review of Genetics, 43(1), pp. 389–410. Available at: 10.1146/annurev-genet-102108-134815.

33. Hoff, K.J. et al. (2016) ‘BRAKER1: Unsupervised RNA-Seq-Based Genome Annotation with GeneMark-ET and AUGUSTUS’, *Bioinformatics (Oxford*, England), 32(5), pp. 767–769. Available at: 10.1093/bioinformatics/btv661.

34. Hust, J. et al. (2018) ‘The Fat-Dachsous signaling pathway regulates growth of horns in Trypoxylus dichotomus, but does not affect horn allometry’, Journal of Insect Physiology, 105, pp. 85–94. Available at: 10.1016/j.jinsphys.2018.01.006.

35. Irvine, K.D. and Harvey, K.F. (2015) ‘Control of organ growth by patterning and hippo signaling in Drosophila’, Cold Spring Harbor Perspectives in Biology, 7(6), p. a019224. Available at: 10.1101/cshperspect.a019224.

36. Järvinen, O. and Vepsäläinen, K. (1976) ‘Wing dimorphism as an adaptive strategy in water-striders (Gerris)’, Hereditas, 84, pp. 61–68.

37. Kaitala, A. and Huldén, L. (1990) ‘Significance of spring migration and flexibility in flight-muscle histolysis in waterstriders (Heteroptera, Gerridae)’, Ecological Entomology, 15(4), pp. 409–418. Available at: 10.1111/j.1365-2311.1990.tb00824.x.

38. Kriventseva, E.V. et al. (2019) ‘OrthoDB v10: sampling the diversity of animal, plant, fungal, protist, bacterial and viral genomes for evolutionary and functional annotations of orthologs’, Nucleic Acids Research, 47(D1), pp. D807–D811. Available at: 10.1093/nar/gky1053.

39. Langfelder, P. and Horvath, S. (2008) ‘WGCNA: an R package for weighted correlation network analysis’, BMC Bioinformatics, 9(1), p. 559. Available at: 10.1186/1471-2105-9-559.

40. Li, H. (2013) ‘Aligning sequence reads, clone sequences and assembly contigs with BWA-MEM’. arXiv. Available at: 10.48550/arXiv.1303.3997.

41. Liao, Y., Smyth, G.K. and Shi, W. (2014) ‘featureCounts: an efficient general purpose program for assigning sequence reads to genomic features’, Bioinformatics, 30(7), pp. 923–930. Available at: 10.1093/bioinformatics/btt656.

42. Lin, X. et al. (2016) ‘JNK signaling mediates wing form polymorphism in brown planthoppers (Nilaparvata lugens)’, Insect Biochemistry and Molecular Biology, 73, pp. 55–61. Available at: 10.1016/j.ibmb.2016.04.005.

43. Lin, X. et al. (2018) ‘Host quality induces phenotypic plasticity in a wing polyphenic insect’, Proceedings of the National Academy of Sciences, 115(29), pp. 7563–7568. Available at: 10.1073/pnas.1721473115.

44. Lobbia, S., Niitsu, S. and Fujiwara, H. (2003) ‘Female-specific wing degeneration caused by ecdysteroid in the Tussock Moth, Orgyia recens: Hormonal and developmental regulation of sexual dimorphism’, Journal of Insect Science, 3, p. 11.

45. Manni, M. et al. (2021) ‘BUSCO Update: Novel and Streamlined Workflows along with Broader and Deeper Phylogenetic Coverage for Scoring of Eukaryotic, Prokaryotic, and Viral Genomes’, Molecular Biology and Evolution, 38(10), pp. 4647–4654. Available at: 10.1093/molbev/msab199.

46. Mao, Y. et al. (2006) ‘Dachs: an unconventional myosin that functions downstream of Fat to regulate growth, affinity and gene expression in Drosophila’, Development, 133(13), pp. 2539– 2551. Available at: 10.1242/dev.02427.

47. Martin, M. (2011) ‘Cutadapt removes adapter sequences from high-throughput sequencing reads’, EMBnet.journal, 17(1), pp. 10–12. Available at: 10.14806/ej.17.1.200.

48. Matakatsu, H. and Blair, S.S. (2006) ‘Separating the adhesive and signaling functions of the Fat and Dachsous protocadherins’, Development, 133(12), pp. 2315–2324. Available at: 10.1242/dev.02401.

49. McCulloch, G.A. et al. (2019) ‘Comparative transcriptomic analysis of a wing-dimorphic stonefly reveals candidate wing loss genes’, EvoDevo, 10(1), p. 21. Available at: 10.1186/s13227-019-0135-4.

50. Mirth, C.K. and Shingleton, A.W. (2019) ‘Coordinating Development: How Do Animals Integrate Plastic and Robust Developmental Processes?’, Frontiers in Cell and Developmental Biology, 7, p. 8. Available at: 10.3389/fcell.2019.00008.

51. Misra, J.R. and Irvine, K.D. (2018) ‘The Hippo Signaling Network and Its Biological Functions’, Annual Review of Genetics, 52(1), pp. 65–87. Available at: 10.1146/annurev-genet-120417-031621.

52. Neumann, C.J. and Cohen, S.M. (1996) ‘A hierarchy of cross-regulation involving Notch, wingless, vestigial and cut organizes the dorsal/ventral axis of the Drosophila wing’, *Development (Cambridge*, England), 122(11), pp. 3477–3485. Available at: 10.1242/dev.122.11.3477.

53. Nijhout, H.F. (2003a) ‘Development and evolution of adaptive polyphenisms’, Evolution & Development, 5(1), pp. 9–18. Available at: 10.1046/j.1525-142X.2003.03003.x.

54. Nijhout, H.F. (2003b) ‘The control of body size in insects’, Developmental Biology, 261(1), pp. 1–9. Available at: 10.1016/S0012-1606(03)00276-8.

55. Nijhout, H.F. (2009) ‘Photoperiodism in Insects: Effects on Morphology’, in R.J. Nelson, D.L. Denlinger, and D.E. Somers (eds) Photoperiodism: The Biological Calendar. Oxford University Press, p. 0. Available at: 10.1093/acprof:oso/9780195335903.003.0013.

56. Nijhout, H.F. and Callier, V. (2015) ‘Developmental Mechanisms of Body Size and Wing-Body Scaling in Insects’, Annual Review of Entomology, 60(1), pp. 141–156. Available at: 10.1146/annurev-ento-010814-020841.

57. Nijhout, H.F., Laub, E. and Grunert, L.W. (2018) ‘Hormonal control of growth in the wing imaginal disks of Junonia coenia: the relative contributions of insulin and ecdysone’, *Development (Cambridge*, England), 145(6), p. dev160101. Available at: 10.1242/dev.160101.

58. Nijhout, H.F. and McKenna, K.Z. (2018) ‘The distinct roles of insulin signaling in polyphenic development’, Current Opinion in Insect Science, 25, pp. 58–64. Available at: 10.1016/j.cois.2017.11.011.

59. Nogueira Alves, A., et al. (2022) ‘Ecdysone coordinates plastic growth with robust pattern in the developing wing’, eLife. Edited by L.M. Riddiford, K.V. Raghavan, and L.M. Riddiford, 11, p. e72666. Available at: 10.7554/eLife.72666.

60. Ohde, T., Mito, T. and Niimi, T. (2022) ‘A hemimetabolous wing development suggests the wing origin from lateral tergum of a wingless ancestor’, Nature Communications, 13(1), p. 979. Available at: 10.1038/s41467-022-28624-x.

61. Parker, J. and Struhl, G. (2020) ‘Control of Drosophila wing size by morphogen range and hormonal gating’, Proceedings of the National Academy of Sciences, 117(50), pp. 31935–31944. Available at: 10.1073/pnas.2018196117.

62. Partridge, L. et al. (1994) ‘EVOLUTION AND DEVELOPMENT OF BODY SIZE AND CELL SIZE IN DROSOPHILA MELANOGASTER IN RESPONSE TO TEMPERATURE’, Evolution; International Journal of Organic Evolution, 48(4), pp. 1269–1276. Available at: 10.1111/j.1558-5646.1994.tb05311.x.

63. Perez-Mockus, G. et al. (2023) ‘The Drosophila ecdysone receptor promotes or suppresses proliferation according to ligand level’, Developmental Cell, 58(20), pp. 2128–2139.e4. Available at: 10.1016/j.devcel.2023.08.032.

64. Poelchau, M. et al. (2015) ‘The i5k Workspace@NAL—enabling genomic data access, visualization and curation of arthropod genomes’, Nucleic Acids Research, 43(D1), pp. D714– D719. Available at: 10.1093/nar/gku983.

65. R Core Team (2022) R: A language and environment for statistical computing., R Foundation for Statistical Computing, Vienna, Austria*. URL* https://www.R-project.org/. Available at: https://www.r-project.org/ (Accessed: 26 March 2024).

66. Ray, R.P. et al. (2015) ‘Patterned Anchorage to the Apical Extracellular Matrix Defines Tissue Shape in the Developing Appendages of Drosophila’, Developmental Cell, 34(3), pp. 310–322. Available at: 10.1016/j.devcel.2015.06.019.

67. Robertson, F.W. (1963) ‘The ecological genetics of growth in Drosophila 6. The genetic correlation between the duration of the larval period and body size in relation to larval diet.’, Genetics Research, 4(1), pp. 74–92. Available at: 10.1017/S001667230000344X.

68. Robinson, M.D., McCarthy, D.J. and Smyth, G.K. (2010) ‘edgeR: a Bioconductor package for differential expression analysis of digital gene expression data’, Bioinformatics, 26(1), pp. 139–140. Available at: 10.1093/bioinformatics/btp616.

69. Roff, D.A. (1984) ‘The cost of being able to fly: a study of wing polymorphism in two species of crickets’, Oecologia, 63(1), pp. 30–37. Available at: 10.1007/BF00379781.

70. Roff, D.A. (1986) ‘The evolution of wing dimorphism in insects’, Evolution; International Journal of Organic Evolution, 40(5), pp. 1009–1020. Available at: 10.1111/j.1558-5646.1986.tb00568.x.

71. Rogulja, D., Rauskolb, C. and Irvine, K.D. (2008) ‘Morphogen Control of Wing Growth through the Fat Signaling Pathway’, Developmental Cell, 15(2), pp. 309–321. Available at: 10.1016/j.devcel.2008.06.003.

72. Rountree, D.B. and Nijhout, H.F. (1995) ‘Hormonal control of a seasonal polyphenism in Precis coenia (Lepidoptera: Nymphalidae)’, Journal of Insect Physiology, 41(11), pp. 987–992. Available at: 10.1016/0022-1910(95)00046-W.

73. Slater, G.S.C. and Birney, E. (2005) ‘Automated generation of heuristics for biological sequence comparison’, BMC Bioinformatics, 6(1), p. 31. Available at: 10.1186/1471-2105-6-31.

74. Smit, A., Hubley, R. and Green, P. (2013) ‘RepeatMasker Open-4.0’. Available at: https://www.repeatmasker.org (Accessed: 13 December 2022).

75. Smýkal, V. et al. (2020) ‘Complex Evolution of Insect Insulin Receptors and Homologous Decoy Receptors, and Functional Significance of Their Multiplicity’, Molecular Biology and Evolution, 37(6), pp. 1775–1789. Available at: 10.1093/molbev/msaa048.

76. Spence, J.R. (1989) ‘The habitat templet and life history strategies of pond skaters (Heteroptera: Gerridae): reproductive potential, phenology, and wing dimorphism’, Canadian Journal of Zoology, 67(10), pp. 2432–2447. Available at: 10.1139/z89-344.

77. Stanke, M. et al. (2008) ‘Using native and syntenically mapped cDNA alignments to improve de novo gene finding’, Bioinformatics, 24(5), pp. 637–644. Available at: 10.1093/bioinformatics/btn013.

78. Steel, C.G.H. and Vafopoulou, X. (2006) ‘Circadian orchestration of developmental hormones in the insect, Rhodnius prolixus’, Comparative Biochemistry and Physiology Part A: Molecular & Integrative Physiology, 144(3), pp. 351–364. Available at: 10.1016/j.cbpa.2006.02.018.

79. Strassburger, K. et al. (2021) ‘Ecdysone regulates Drosophila wing disc size via a TORC1 dependent mechanism’, Nature Communications, 12(1), p. 6684. Available at: 10.1038/s41467-021-26780-0.

80. Tanimoto, H. et al. (2000) ‘Hedgehog creates a gradient of DPP activity in Drosophila wing imaginal discs’, Molecular Cell, 5(1), pp. 59–71. Available at: 10.1016/s1097-2765(00)80403-7.

81. Tripathi, B.K. and Irvine, K.D. (2022) ‘The wing imaginal disc’, Genetics, 220(4), p. iyac020. Available at: 10.1093/genetics/iyac020.

82. Uyehara, C.M., Leatham-Jensen, M. and McKay, D.J. (2022) ‘Opportunistic binding of EcR to open chromatin drives tissue-specific developmental responses’, Proceedings of the National Academy of Sciences, 119(40), p. e2208935119. Available at: 10.1073/pnas.2208935119.

83. Uyehara, C.M. and McKay, D.J. (2019) ‘Direct and widespread role for the nuclear receptor EcR in mediating the response to ecdysone in Drosophila’, Proceedings of the National Academy of Sciences, 116(20), pp. 9893–9902. Available at: 10.1073/pnas.1900343116.

84. Vellichirammal, N., Madayiputhiya, N. and Brisson, J.A. (2016) ‘The genome-wide transcriptional response underlying the pea aphid wing polyphenism’, Mol Ecol [Preprint]. Available at: 10.1111/mec.13749.

85. Vellichirammal, N.N. et al. (2017) ‘Ecdysone signaling underlies the pea aphid transgenerational wing polyphenism’, Proc Natl Acad Sci U S A [Preprint]. Available at: 10.1073/pnas.1617640114.

86. Vepsäläinen, K. (1978) ‘Wing Dimorphism and Diapause in Gerris: Determination and Adaptive Significance’, in, pp. 218–253. Available at: 10.1007/978-1-4615-6941-1_10.

87. Verghese, S., Bedi, S. and Kango-Singh, M. (2012) ‘Hippo signalling controls Dronc activity to regulate organ size in Drosophila’, Cell Death & Differentiation, 19(10), pp. 1664–1676. Available at: 10.1038/cdd.2012.48.

88. Walker, B.J. et al. (2014) ‘Pilon: An Integrated Tool for Comprehensive Microbial Variant Detection and Genome Assembly Improvement’, PLOS ONE, 9(11), p. e112963. Available at: 10.1371/journal.pone.0112963.

89. Willecke, M. et al. (2008) ‘Boundaries of Dachsous Cadherin activity modulate the Hippo signaling pathway to induce cell proliferation’, Proceedings of the National Academy of Sciences, 105(39), pp. 14897–14902. Available at: 10.1073/pnas.0805201105.

90. Xu, H.J. et al. (2015) ‘Two insulin receptors determine alternative wing morphs in planthoppers’, Nature, 519(7544), pp. 464–7. Available at: 10.1038/nature14286.

91. Xu, H.J. and Zhang, C.X. (2017) ‘Insulin receptors and wing dimorphism in rice planthoppers’, Philos Trans R Soc Lond B Biol Sci, 372(1713). Available at: 10.1098/rstb.2015.0489.

92. Ye, X. et al. (2019) ‘miR-34 modulates wing polyphenism in planthopper’, PLOS Genetics, 15(6), p. e1008235. Available at: 10.1371/journal.pgen.1008235.

93. Zera, A.J. (1984) ‘Differences in Survivorship, Development Rate and Fertility Between the Longwinged and Wingless Morphs of the Waterstrider, Limnoporus Canaliculatus’, Evolution, 38(5), pp. 1023–1032. Available at: 10.1111/j.1558-5646.1984.tb00372.x.

94. Zera, A.J. and Denno, R.F. (1997) ‘Physiology and Ecology of Dispersal Polymorphism in Insects’, Annual Review of Entomology, 42(1), pp. 207–230. Available at: 10.1146/annurev.ento.42.1.207.

95. Zera, A.J. and Larsen, A. (2001) ‘The metabolic basis of life history variation: genetic and phenotypic differences in lipid reserves among life history morphs of the wing-polymorphic cricket, Gryllus firmus’, Journal of Insect Physiology, 47(10), pp. 1147–1160. Available at: 10.1016/s0022-1910(01)00096-8.

96. Zhang, C. et al. (2015) ‘The Ecdysone Receptor Coactivator Taiman Links Yorkie to Transcriptional Control of Germline Stem Cell Factors in Somatic Tissue’, Developmental Cell, 34(2), pp. 168–180. Available at: 10.1016/j.devcel.2015.05.010.

97. Zhang, C.-X., Brisson, J.A. and Xu, H.-J. (2019) ‘Molecular Mechanisms of Wing Polymorphism in Insects’, Annual Review of Entomology, 64(1), pp. 297–314. Available at: 10.1146/annurev-ento-011118-112448.

98. Zhang, J.-L. et al. (2021) ‘Vestigial mediates the effect of insulin signaling pathway on wing-morph switching in planthoppers’, PLOS Genetics, 17(2), p. e1009312. Available at: 10.1371/journal.pgen.1009312.

99. Zhang, J.-L. et al. (2022) ‘The transcription factor Zfh1 acts as a wing-morph switch in planthoppers’, Nature Communications, 13(1), p. 5670. Available at: 10.1038/s41467-022-33422-6.

100. Atkinson, D. 1994. ‘Temperature and Organism Size—A Biological Law for Ectotherms?’ In Advances in Ecological Research, edited by M. Begon and A. H. Fitter, 25:1–58. Academic Press. 10.1016/S0065-2504(08)60212-3.

101. Bakker, K. 1959. ‘Feeding Period, Growth, and Pupation in Larvae Ofdrosophila Melanogaster’. Entomologia Experimentalis et Applicata 2 (3): 171–86. 10.1111/j.1570-7458.1959.tb00432.x.

102. Brogiolo, Walter, Hugo Stocker, Tomoatsu Ikeya, Felix Rintelen, Rafael Fernandez, and Ernst Hafen. 2001. ‘An Evolutionarily Conserved Function of the Drosophila Insulin Receptor and Insulin-like Peptides in Growth Control’. Current Biology 11 (4): 213–21. 10.1016/S0960-9822(01)00068-9.

103. Burg, Karin R. L. van der, James J. Lewis, Benjamin J. Brack, Richard A. Fandino, Anyi Mazo-Vargas, and Robert D. Reed. 2020. ‘Genomic Architecture of a Genetically Assimilated Seasonal Color Pattern’. Science 370 (6517): 721–25. 10.1126/science.aaz3017.

104. Dobin, Alexander, Carrie A. Davis, Felix Schlesinger, Jorg Drenkow, Chris Zaleski, Sonali Jha, Philippe Batut, Mark Chaisson, and Thomas R. Gingeras. 2013. ‘STAR: Ultrafast Universal RNA-Seq Aligner’. Bioinformatics 29 (1): 15–21. 10.1093/bioinformatics/bts635.

105. Fawcett, Meghan M., Mary C. Parks, Alice E. Tibbetts, Jane S. Swart, Elizabeth M. Richards, Juan Camilo Vanegas, Meredith Cenzer, et al. 2018. ‘Manipulation of Insulin Signaling Phenocopies Evolution of a Host-Associated Polyphenism’. Nature Communications 9 (1): 1699. 10.1038/s41467-018-04102-1.

106. Gokhale, Rewatee H., Takashi Hayashi, Christopher D. Mirque, and Alexander W. Shingleton. 2016. ‘Intra-Organ Growth Coordination in Drosophila Is Mediated by Systemic Ecdysone Signaling’. Developmental Biology 418 (1): 135–45. 10.1016/j.ydbio.2016.07.016.

107. Gotoh, Hiroki, James A. Hust, Toru Miura, Teruyuki Niimi, Douglas J. Emlen, and Laura C. Lavine. 2015. ‘The Fat/Hippo Signaling Pathway Links within-Disc Morphogen Patterning to Whole-Animal Signals during Phenotypically Plastic Growth in Insects’. Developmental Dynamics 244 (9): 1039–45. 10.1002/dvdy.24296.

108. Gudmunds, Erik, Shrinath Narayanan, Elise Lachivier, Marion Duchemin, Abderrahman Khila, and Arild Husby. 2022. ‘Photoperiod Controls Wing Polyphenism in a Water Strider Independently of Insulin Receptor Signalling’. Proceedings of the Royal Society B: Biological Sciences 289 (1973): 20212764. 10.1098/rspb.2021.2764.

109. Hayes, Abigail M., Mark D. Lavine, Hiroki Gotoh, Xinda Lin, and Laura Corley Lavine. 2019. ‘Mechanisms Regulating Phenotypic Plasticity in Wing Polyphenic Insects’. In Advances in Insect Physiology, edited by Russell Jurenka, 56:43–72. Academic Press. 10.1016/bs.aiip.2019.01.005.

110. Herboso, Leire, Marisa M. Oliveira, Ana Talamillo, Coralia Pérez, Monika González, David Martín, James D. Sutherland, Alexander W. Shingleton, Christen K. Mirth, and Rosa Barrio. 2015. ‘Ecdysone Promotes Growth of Imaginal Discs through the Regulation of Thor in D. Melanogaster’. Scientific Reports 5 (1): 12383. 10.1038/srep12383.

111. Hietakangas, Ville, and Stephen M. Cohen. 2009. ‘Regulation of Tissue Growth through Nutrient Sensing’. Annual Review of Genetics 43 (1): 389–410. 10.1146/annurev-genet-102108-134815.

112. Liao, Yang, Gordon K. Smyth, and Wei Shi. 2014. ‘featureCounts: An Efficient General Purpose Program for Assigning Sequence Reads to Genomic Features’. Bioinformatics 30 (7): 923–30. 10.1093/bioinformatics/btt656.

113. Lin, Xinda, Yili Xu, Jianru Jiang, Mark Lavine, and Laura Corley Lavine. 2018. ‘Host Quality Induces Phenotypic Plasticity in a Wing Polyphenic Insect’. Proceedings of the National Academy of Sciences 115 (29): 7563–68. 10.1073/pnas.1721473115.

114. Lin, Xinda, Yili Xu, Yun Yao, Bo Wang, Mark D. Lavine, and Laura Corley Lavine. 2016. ‘JNK Signaling Mediates Wing Form Polymorphism in Brown Planthoppers (Nilaparvata Lugens)’. Insect Biochemistry and Molecular Biology 73 (June): 55–61. 10.1016/j.ibmb.2016.04.005.

115. Martin, Marcel. 2011. ‘Cutadapt Removes Adapter Sequences from High-Throughput Sequencing Reads’. EMBnet.Journal 17 (1): 10–12. 10.14806/ej.17.1.200.

116. Mirth, Christen K., and Alexander W. Shingleton. 2019. ‘Coordinating Development: How Do Animals Integrate Plastic and Robust Developmental Processes?’ Frontiers in Cell and Developmental Biology 7: 8. 10.3389/fcell.2019.00008.

117. Nijhout, H. F. 2003. ‘The Control of Body Size in Insects’. Developmental Biology 261 (1): 1–9. 10.1016/S0012-1606(03)00276-8.

118. Nijhout, H. Frederik. 2003. ‘Development and Evolution of Adaptive Polyphenisms’. Evolution & Development 5 (1): 9–18. 10.1046/j.1525-142X.2003.03003.x.

119. Nijhout, H. Frederik, and Viviane Callier. 2015. ‘Developmental Mechanisms of Body Size and Wing-Body Scaling in Insects’. Annual Review of Entomology 60 (1): 141–56. 10.1146/annurev-ento-010814-020841.

120. Nijhout, H. Frederik, Emily Laub, and Laura W. Grunert. 2018. ‘Hormonal Control of Growth in the Wing Imaginal Disks of Junonia Coenia: The Relative Contributions of Insulin and Ecdysone’. *Development (Cambridge*, England) 145 (6): dev160101. 10.1242/dev.160101.

121. Nijhout, H Frederik, and Kenneth Z McKenna. 2018. ‘The Distinct Roles of Insulin Signaling in Polyphenic Development’. *Current Opinion in Insect Science*, Insect genomics * Development and regulation, 25 (February): 58–64. 10.1016/j.cois.2017.11.011.

122. Nogueira Alves, André, Marisa Mateus Oliveira, Takashi Koyama, Alexander Shingleton, and Christen Kerry Mirth. 2022. ‘Ecdysone Coordinates Plastic Growth with Robust Pattern in the Developing Wing’. Edited by Lynn M Riddiford, K Vijay Raghavan, and Lynn M Riddiford. eLife 11 (March): e72666. 10.7554/eLife.72666.

123. Parker, Joseph, and Gary Struhl. 2020. ‘Control of Drosophila Wing Size by Morphogen Range and Hormonal Gating’. Proceedings of the National Academy of Sciences 117 (50): 31935–44. 10.1073/pnas.2018196117.

124. Partridge, Linda, Brian Barrie, Kevin Fowler, and Vernon French. 1994. ‘EVOLUTION AND DEVELOPMENT OF BODY SIZE AND CELL SIZE IN DROSOPHILA MELANOGASTER IN RESPONSE TO TEMPERATURE’. Evolution; International Journal of Organic Evolution 48 (4): 1269–76. 10.1111/j.1558-5646.1994.tb05311.x.

125. Perez-Mockus, Gantas, Luca Cocconi, Cyrille Alexandre, Birgit Aerne, Guillaume Salbreux, and Jean-Paul Vincent. 2023. ‘The Drosophila Ecdysone Receptor Promotes or Suppresses Proliferation According to Ligand Level’. Developmental Cell 58 (20): 2128–2139.e4. 10.1016/j.devcel.2023.08.032.

126. Robertson, Forbes W. 1963. ‘The Ecological Genetics of Growth in Drosophila 6. The Genetic Correlation between the Duration of the Larval Period and Body Size in Relation to Larval Diet.’ Genetics Research 4 (1): 74–92. 10.1017/S001667230000344X.

127. Rountree, D. B., and H. F. Nijhout. 1995. ‘Hormonal Control of a Seasonal Polyphenism in Precis Coenia (Lepidoptera: Nymphalidae)’. Journal of Insect Physiology 41 (11): 987–92. 10.1016/0022-1910(95)00046-W.

128. Smýkal, Vlastimil, Martin Pivarči, Jan Provazník, Olga Bazalová, Pavel Jedlička, Ondřej Lukšan, Aleš Horák, et al. 2020. ‘Complex Evolution of Insect Insulin Receptors and Homologous Decoy Receptors, and Functional Significance of Their Multiplicity’. Molecular Biology and Evolution 37 (6): 1775–89. 10.1093/molbev/msaa048.

129. Spence, John R. 1989. ‘The Habitat Templet and Life History Strategies of Pond Skaters (Heteroptera: Gerridae): Reproductive Potential, Phenology, and Wing Dimorphism’. Canadian Journal of Zoology 67 (10): 2432–47. 10.1139/z89-344.

130. Strassburger, Katrin, Marilena Lutz, Sandra Müller, and Aurelio A. Teleman. 2021. ‘Ecdysone Regulates Drosophila Wing Disc Size via a TORC1 Dependent Mechanism’. Nature Communications 12 (1): 6684. 10.1038/s41467-021-26780-0.

131. Tripathi, Bipin Kumar, and Kenneth D Irvine. 2022. ‘The Wing Imaginal Disc’. Genetics 220 (4): iyac020. 10.1093/genetics/iyac020.

132. Vellichirammal, N., N. Madayiputhiya, and J. A. Brisson. 2016. ‘The Genome-Wide Transcriptional Response Underlying the Pea Aphid Wing Polyphenism’. *Mol Ecol*, July. 10.1111/mec.13749.

133. Vellichirammal, N. N., P. Gupta, T. A. Hall, and J. A. Brisson. 2017. ‘Ecdysone Signaling Underlies the Pea Aphid Transgenerational Wing Polyphenism’. *Proc Natl Acad Sci U S A*, January. 10.1073/pnas.1617640114.

134. Xu, H. J., J. Xue, B. Lu, X. C. Zhang, J. C. Zhuo, S. F. He, X. F. Ma, et al. 2015. ‘Two Insulin Receptors Determine Alternative Wing Morphs in Planthoppers’. Nature 519 (7544): 464–67. 10.1038/nature14286.

135. Zhang, Jin-Li, Sheng-Jie Fu, Sun-Jie Chen, Hao-Hao Chen, Yi-Lai Liu, Xin-Yang Liu, and Hai-Jun Xu. 2021. ‘Vestigial Mediates the Effect of Insulin Signaling Pathway on Wing-Morph Switching in Planthoppers’. PLOS Genetics 17 (2): e1009312. 10.1371/journal.pgen.1009312.

