## Supplemental Material for "The Fat/Hippo pathway drives photoperiod-induced wing length polyphenism"

### Supplemental Files

### Supplementary Files

**Supplementary File 1. Results of the differential expression analysis at i4L**
**Supplementary File 2. Results of the differential expression analysis at i5L**
**Supplementary File 3. Results of the GO Term enrichment analysis at i4L**
**Supplementary File 4. Results of the GO Term enrichment analysis at i5L**
**Supplementary File 5. Results of the GO Term enrichment analysis for Module 1**
**Supplementary File 6. Results of the GO Term enrichment analysis for Module 2**
**Supplementary File 7. Results of the GO Term enrichment analysis for Module 4**
**Supplementary File 8. Results of the GO Term enrichment analysis for Module 5**

**Table S1. Developmental duration in 12L:12D and 18L:6D.**

| Mean (days) | SD | Stage | Photoperiod |
|---|---|---|---|
| 3,1 | 0,3 | Instar 3 | 12L:12D |
| 3,6 | 0,5 | Instar 4 | 12L:12D |
| 5,8 | 0,7 | Instar 5 | 12L:12D |
| 3,1 | 0,3 | Instar 3 | 18L:6D |
| 3,3 | 0,5 | Instar 4 | 18L:6D |
| 4,7 | 0,5 | Instar 5 | 18L:6D |

**Table S2. Number of total reads and mapping percentage for each RNA-seq library.**

| Library | nReads | %Mapped | Discarded? |
|---|---|---|---|
| i4E_18_2 | 28169836 | 68,34 | No |
| i4E_18_3 | 95871243 | 69,98 | No |
| i4E_18_4 | 85556299 | 65,71 | No |
| i4E_18_5 | 79254911 | 69,01 | No |
| i4E_18_6 | 79290348 | 67,10 | No |
| i4E_18_7 | 33577301 | 63,56 | No |
| i4L_18_1 | 72213722 | 72,93 | **Yes** |
| i4L_18_2 | 72606303 | 70,51 | No |
| i4L_18_5 | 71137632 | 70,05 | No |
| i4L_18_6 | 75403194 | 69,78 | No |
| i4L_18_7 | 78047003 | 65,42 | No |
| i5E_18_1 | 76754978 | 67,22 | No |
| i5E_18_2 | 73494614 | 65,38 | No |
| i5E_18_3 | 63555 | 68,15 | **Yes** |
| i5E_18_4 | 51249278 | 65,90 | No |
| i5E_18_5 | 68790307 | 69,86 | No |
| i5E_18_7 | 78702853 | 70,58 | No |
| i5L_18_1 | 59547856 | 64,01 | **Yes** |
| i5L_18_2 | 14730645 | 41,64 | **Yes** |
| i5L_18_3 | 83209106 | 66,73 | No |
| i5L_18_4 | 32366523 | 70,59 | No |
| i5L_18_5 | 113188 | 65,75 | **Yes** |
| i5L_18_6 | 28642136 | 63,25 | No |
| i4E_12_2 | 74520419 | 68,76 | No |
| i4E_12_3 | 75548669 | 71,63 | No |
| i4E_12_4 | 60331096 | 69,74 | No |
| i4E_12_5 | 64569756 | 71,38 | No |
| i4E_12_6 | 69408608 | 70,62 | No |
| i4E_12_7 | 78193240 | 69,68 | No |
| i4L_12_1 | 30409160 | 76,97 | No |
| i4L_12_3 | 69946164 | 74,18 | **Yes** |
| i4L_12_5 | 94315176 | 74,00 | No |
| i4L_12_6 | 60582463 | 78,50 | No |
| i4L_12_7 | 69732776 | 74,53 | No |
| i4L_12_8 | 69694500 | 68,86 | No |
| i5E_12_1 | 91299202 | 71,59 | No |
| i5E_12_2 | 65743353 | 69,50 | No |
| i5E_12_3 | 28621101 | 72,16 | No |
| i5E_12_4 | 70399189 | 69,25 | No |
| i5E_12_5 | 76884326 | 72,38 | No |
| i5E_12_7 | 73052128 | 75,21 | No |
| i5L_12_1 | 79160905 | 71,12 | No |
| i5L_12_2 | 70618844 | 68,98 | No |
| i5L_12_3 | 75382045 | 73,44 | No |
| i5L_12_4 | 83790014 | 75,42 | **Yes** |
| i5L_12_5 | 95590197 | 76,16 | No |

**Table S3. Gene, gene identifiers, primers and injected approximate dose used in RNAi experiments.** The T7 promotor sequence is displayed in bold letters.

| Gene | Gene ID | Strand | Purpose | Primer sequence |
|---|---|---|---|---|
| *Fat* | gbgene9923 | **Rv** | **dsRNA** | **TAATACGACTCACTATAGGGAGACCAC**TCACTGTGGATGAGAGACTC |
| *Fat* | | **Fw** | | **TAATACGACTCACTATAGGGAGACCAC**CTCAGTGTCGTATTCACTCG |
| *Dachsous* | gbgene4229 | **Rv** | **dsRNA** | **TAATACGACTCACTATAGGGAGACCAC**TTAGGAGGACCGTATTGAGC |
| *Dachsous* | | **Fw** | | **TAATACGACTCACTATAGGGAGACCAC**AGAGATAAAGGACCTGACGG |
| *Yorkie* | gbgene2745 | **Rv** | **dsRNA** | **TAATACGACTCACTATAGGGAGACCAC**AGCTGTAATGAAGGTACGAG |
| *Yorkie* | | **Fw** | | **TAATACGACTCACTATAGGGAGACCAC**ACTTCATGGTTTGACCCAAG |
| *Scalloped* | gbman.gbue005394 | **Rv** | **dsRNA** | **TAATACGACTCACTATAGGGAGACCAC**TGTACCCTGACTGGAAACTC |
| *Scalloped* | | **Fw** | | **TAATACGACTCACTATAGGGAGACCAC**GCTCAGAAAAATCAAGCCTG |
| *Dachs* | gbgene18132 | **Rv** | **dsRNA** | **TAATACGACTCACTATAGGGAGACCAC**GGTCCAAGAAGTAACAGTGG |
| *Dachs* | | **Fw** | | **TAATACGACTCACTATAGGGAGACCAC**TCTTATCCACCTTCCTGGAC |
| *Crumbs* | gbgene5059 | **Rv** | **dsRNA** | **TAATACGACTCACTATAGGGAGACCAC**TTGGGATAAACATTGCAGGG |
| *Crumbs* | | **Fw** | | **TAATACGACTCACTATAGGGAGACCAC**ATCTTTGCAAAAACGATGCC |
| *Dumpy* | gbgene7473 | **Rv** | **dsRNA** | **TAATACGACTCACTATAGGGAGACCAC**GGACATTCACTGTTCTGTCG |
| *Dumpy* | | **Fw** | | **TAATACGACTCACTATAGGGAGACCAC**TCGATGAACGTAACTTGCG |
| *Hr4* | gbgene15190 | **Rv** | **dsRNA** | **TAATACGACTCACTATAGGGAGACCAC**CCAGCTCTGTTGAACCAC |
| *Hr4* | | **Fw** | | **TAATACGACTCACTATAGGGAGACCAC**TCTGTTAGACAACAAAGCTG |
| *L(3)mbn* | gbgene10227 | **Rv** | **dsRNA** | **TAATACGACTCACTATAGGGAGACCAC**TCCGTGGAACTTGTTATGAG |
| *L(3)mbn* | | **Fw** | | **TAATACGACTCACTATAGGGAGACCAC**TAGAGATGGCAGAAGTCGG |
| *Pal1* | gbgene2662 | **Rv** | **dsRNA** | **TAATACGACTCACTATAGGGAGACCAC**ATAATGGCAGCAGCAAATAC |
| *Pal1* | | **Fw** | | **TAATACGACTCACTATAGGGAGACCAC**TGCTTTGACAGTAGTGGAAG |
| *Stubble* | gbgene217 | **Rv** | **dsRNA** | **TAATACGACTCACTATAGGGAGACCAC**CTGAGTTGTTACGTGTAGCC |
| *Stubble* | | **Fw** | | **TAATACGACTCACTATAGGGAGACCAC**CTGAGTTGTTACGTGTAGCC |
| *Su(dx)* | gbgene19356 | **Rv** | **dsRNA** | **TAATACGACTCACTATAGGGAGACCAC**AGGGATCACTGTCTACTGAC |
| *Su(dx)* | | **Fw** | | **TAATACGACTCACTATAGGGAGACCAC**TAGACTGGTGACTCCGTAC |
| *Atx-1* | gbgene16742 | **Rv** | **dsRNA** | **TAATACGACTCACTATAGGGAGACCAC**CTTCTCGAGGGTTTTATCTGG |
| *Atx-1* | | **Fw** | | **TAATACGACTCACTATAGGGAGACCAC**CGACGCCTTATCTTTACCAG |
| *ERR* | gbgene13007 | **Rv** | **dsRNA** | **TAATACGACTCACTATAGGGAGACCAC**CAGACACGTCAACTCTTCTG |
| *ERR* | | **Fw** | | **TAATACGACTCACTATAGGGAGACCAC**TTCGTTGTTACAATCGCCC |
| *Fat* | gbgene9923 | **Rv** | **RT-qPCR** | ACTTGACCAGGCCTTTCAAGA |
| *Fat* | | **Fw** | | CGTCGTACCGGTAAGACTAC |
| *Ds* | gbgene4229 | **Rv** | **RT-qPCR** | GCTTGGACAGTCAAAAGGGC |
| *Ds* | | **Fw** | | TGACAATGTGCCACAGTTCC |
| *Yki* | gbgene2745 | **Rv** | **RT-qPCR** | GCCCTCTGCAGATGAACAGG |
| *Yki* | | **Fw** | | GAACAGGCTACGACCGCAG |
| *RPS26* | gbgene1421 | **Rv** | **RT-qPCR** | AGAAATATCGTTGAA |
| *RPS26* | | **Fw** | | ACAGTAGTGAAGCTT |

**Table S4. WGCNA modules obtained.** The total number of genes clustering in each module as well as the proportion of variation the eigengenes of said modules represent are indicated.

| Module Number | nGenes | Explained Variation |
|---:|---:|---:|
| 1 | 2064 | 0.7509582 |
| 2 | 1404 | 0.6225611 |
| 3 | 921 | 0.6766435 |
| 4 | 251 | 0.6362751 |
| 5 | 205 | 0.7147215 |
| 6 | 151 | 0.7431185 |
| 7 | 138 | 0.8140409 |
| 8 | 101 | 0.8347595 |
| 9 | 99 | 0.6328425 |
| 10 | 52 | 0.6987123 |
| 11 | 51 | 0.8325352 |
| 12 | 41 | 0.6839508 |
| 13 | 41 | 0.7569754 |

**Table S5. Differentially expressed genes in i5L targeted with RNAi in instar four to screen for genes controlling wing morph determination.** By column: 1; Gene ID in the *G. buenoi* annotation, 2; Gene name in *D. melanogaster*, 3; Flybase ID, 4; Log2 fold change, 5; motivation for functional validation.

| GeneID | Gene name | Flybase ID | L2FC | Motivation |
|---|---|---|---|---|
| gbgene9923 | Fat | FBgn0001075 | -0.92 | Involved in Hippo signaling. |
| gbgene4229 | Dachsous | FBgn0284247 | -2.1 | Involved in Hippo signaling. |
| gbgene2745 | Yorkie | FBgn0034970 | -3.3 | Involved in Hippo signaling. |
| gbgene18132 | Dachs | FBgn0262029 | 1.0 | Involved in Hippo signaling. |
| gbgene5059 | Crumbs | FBgn0259685 | -2.3 | Involved in Notch and Hippo signaling. |
| gbgene7473 | Dumpy | FBgn0053196 | -2.0 | Regulation of wing size in *Dme* |
| gbman.GBUE008359-PA.1 | Hr4 | FBgn0264562 | -4.1 | Nuclear receptor transcriptionally induced by ecdysone, transcription factor. |
| gbgene10227 | L(3)mbn | FBgn0002440 | -17.2 | One of the highest L2FC values, tumor suppressor gene. |
| gbgene2662 | Pal1 | FBgn0283510 | -1.2 | Regulation of wing size in interaction with Dpy. |
| gbgene217 | Stubble | FBgn0003319 | -2.9 | Hormone-dependent protease involved in epithelial morphogenesis in wings. |
| gbgene19356 | Su(dx) | FBgn0003557 | -1.4 | E3 ubiquitin-protein ligase involved in Notch signaling. |
| gbgene16742 | Atx-1 | FBgn0029907 | 10.5 | Regulation of transcription, third lowest FDR. |
| gbman.GBUE008299-PA.1 | ERR | FBgn0035849 | -1.3 | Nuclear receptor with DNA binding capacity. |

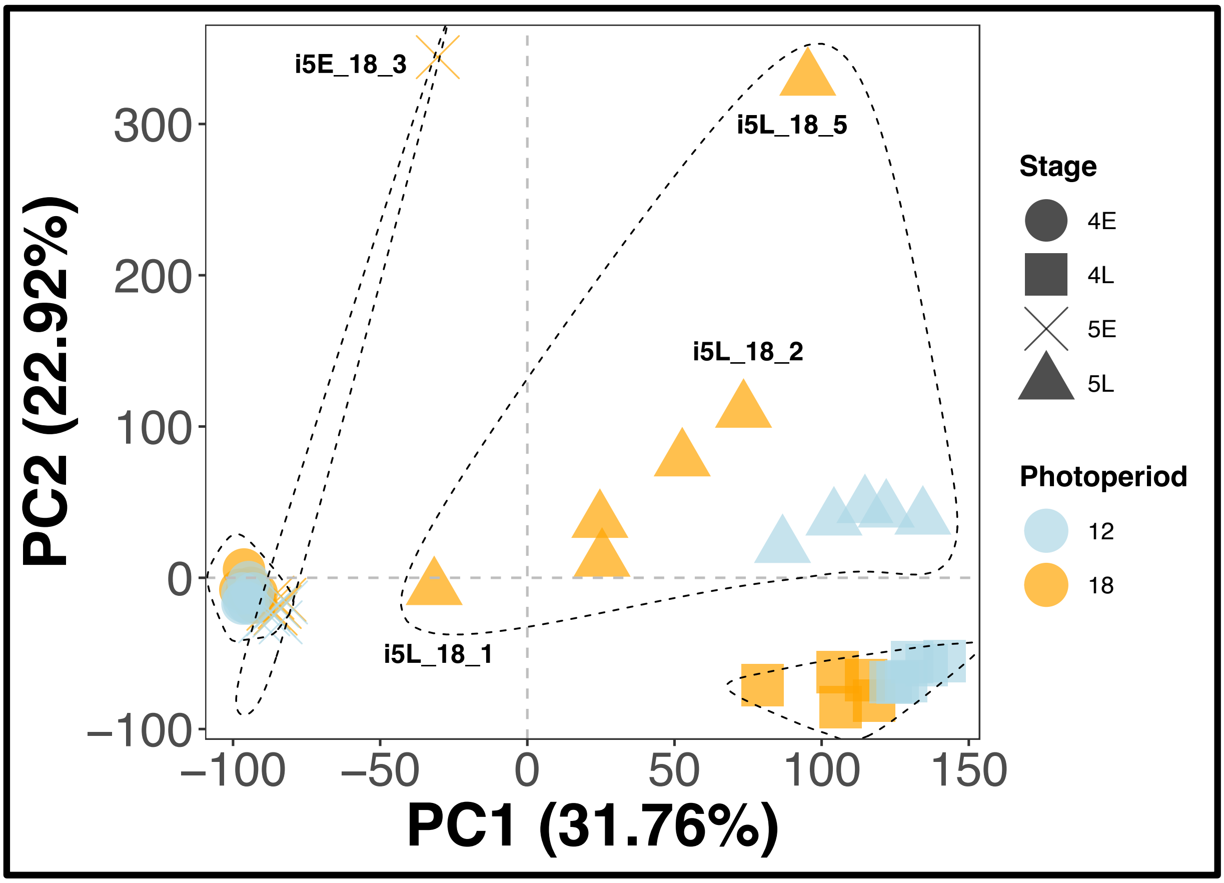

**Figure S1. PCA including all sequencing libraries before filtering.** Indicated libraries were discarded due to poor clustering with the rest of the same treatment (i5E_18_3 and i5L_18_1) or due to low mapped read count (see Supplemental Table S2).

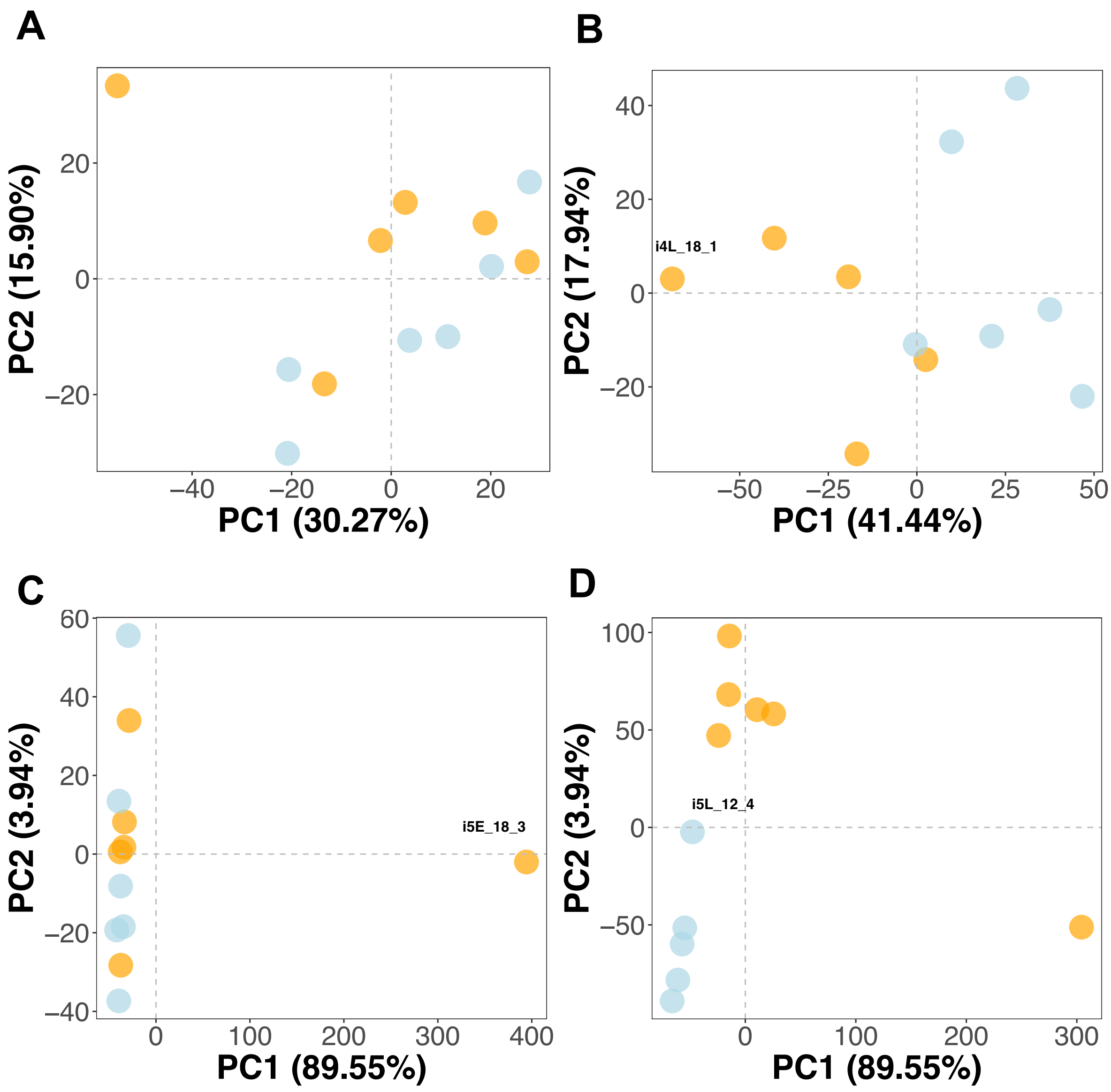

**Figure S2. Stage-specific PCA plots before library filtering.** Libraries indicated with their code were removed based on poor clustering. Blue dots represent libraries at 12L:12D, and orange ones represent libraries at 18L:6D.

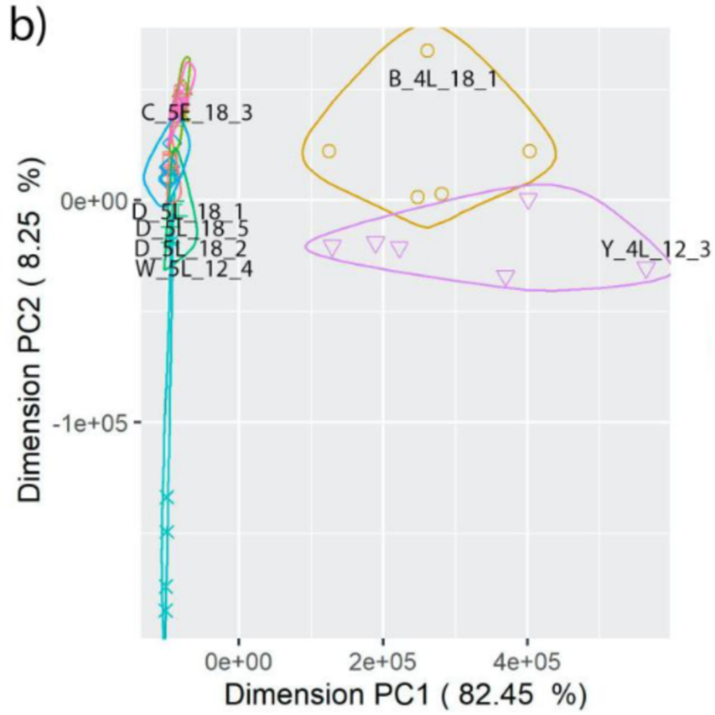

**Figure S3. PCA plot for all prepared libraries using only the data from the 100 most highly expressed genes.** Library i4L_12_3 was eliminated from further analysis due to a large difference on clustering based on PC1 with respect to other libraries in the same treatment.

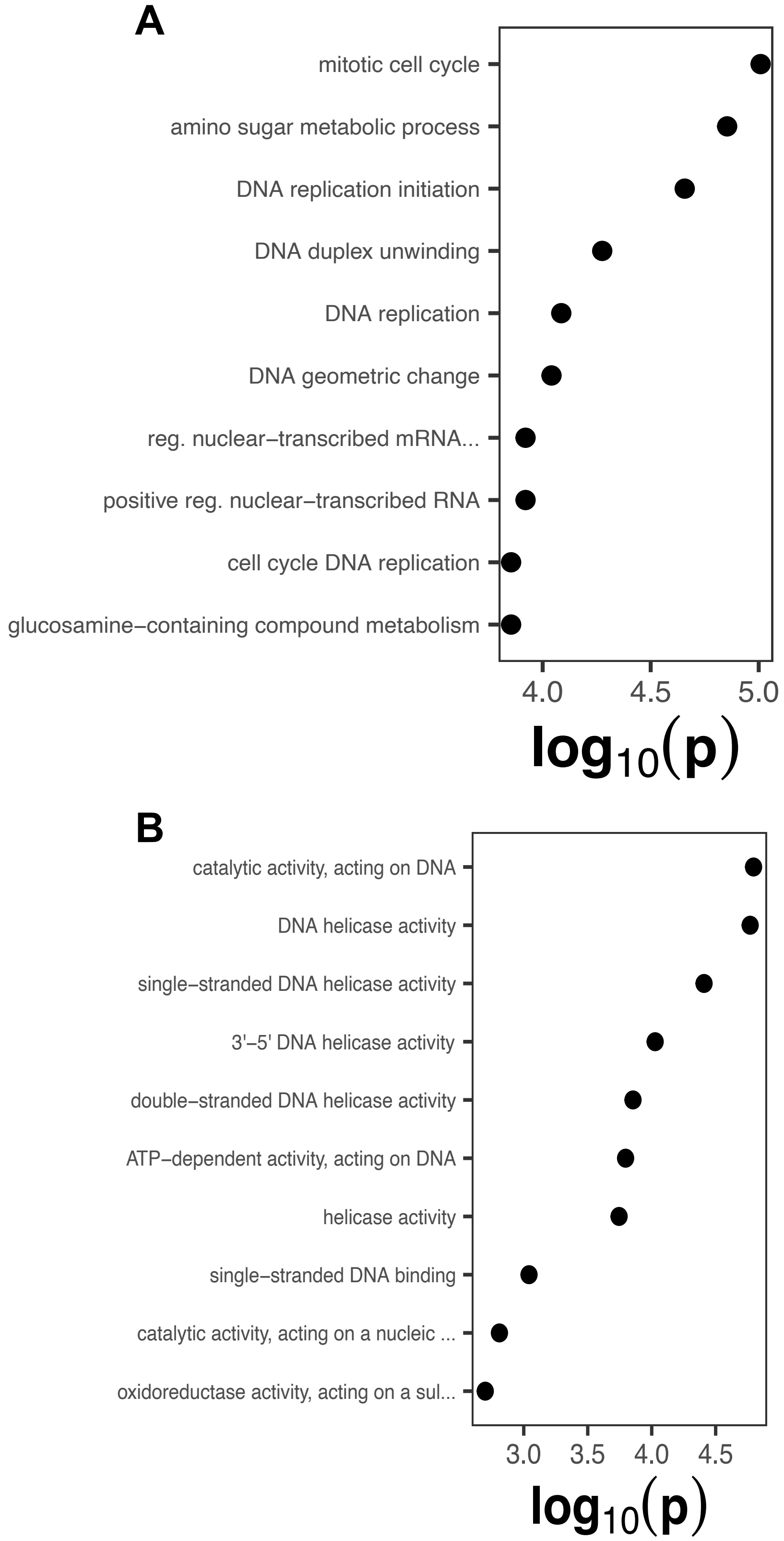

**Figure S4. GO Term enrichment analysis in the comparison between 12L:12D and 18L:6D at i4L, based on the 146 differentially expressed genes.** The 10 most-enriched terms for BP **(A)** and MF **(B)** are shown.

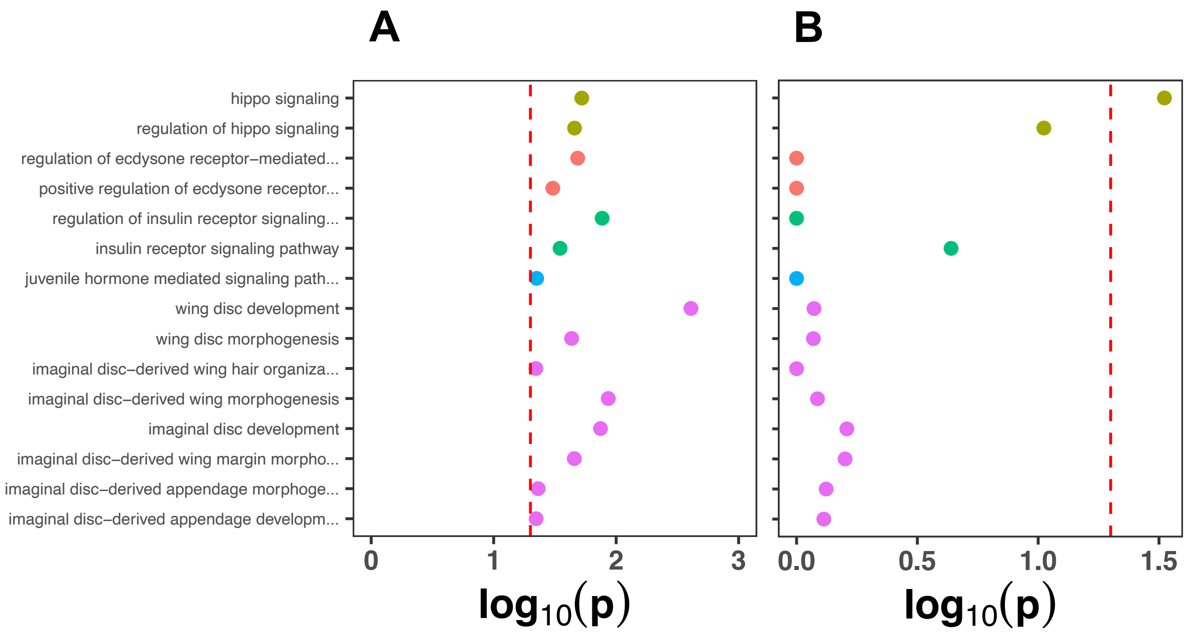

**Figure S5. Results of the GO Term enrichment analysis focusing on candidate pathways responsible for wing formation and development.** All significantly enriched GO Terms in i5L **(A)** were included, associated with the Hippo pathway (yellow), ecdysone signaling (red), insulin signaling (green), juvenile hormone (blue) and imaginal disc and wing formation (purple). For comparison, results for the same GO Terms in i4L **(B)** were included. The vertical dashed line indicates the threshold for significance ($p = 0.05$).

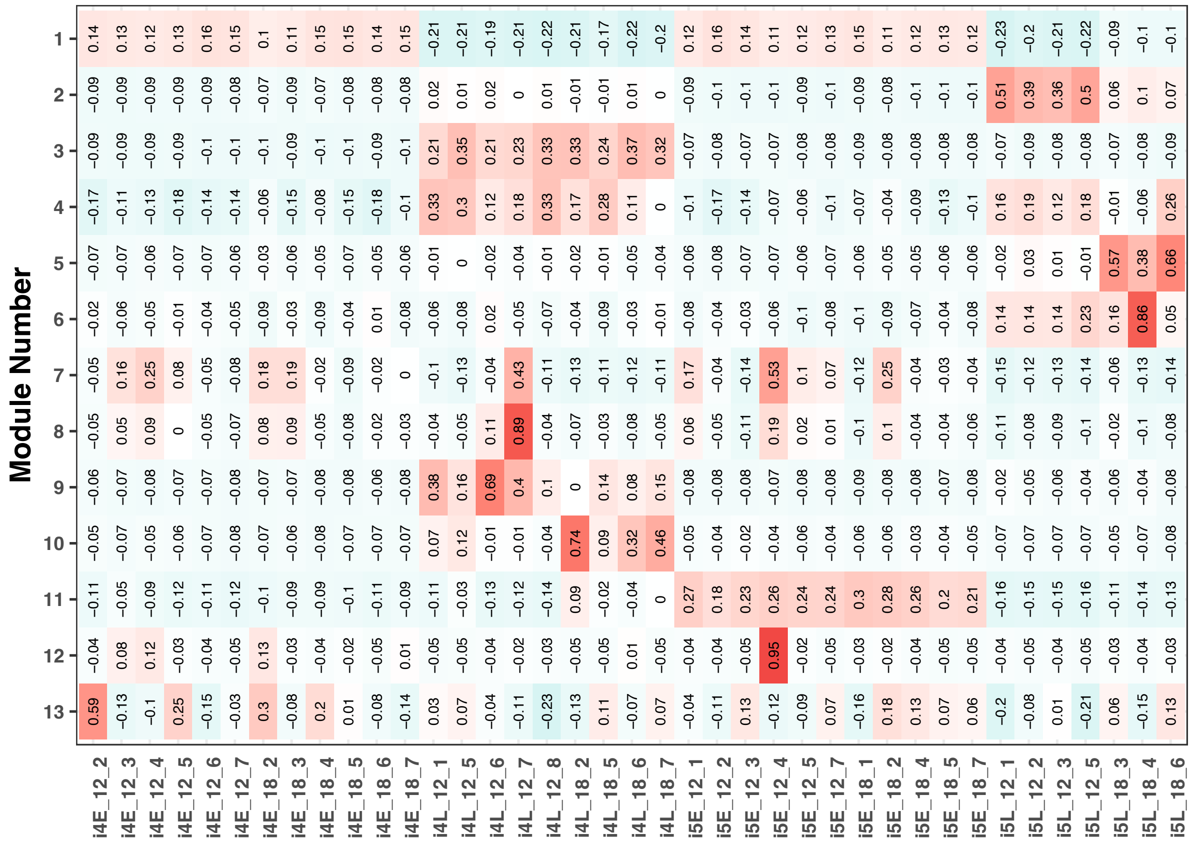

**Figure S6. WGCNA clustering results.** The eigengene correlation with each individual RNA library after filtering is shown.

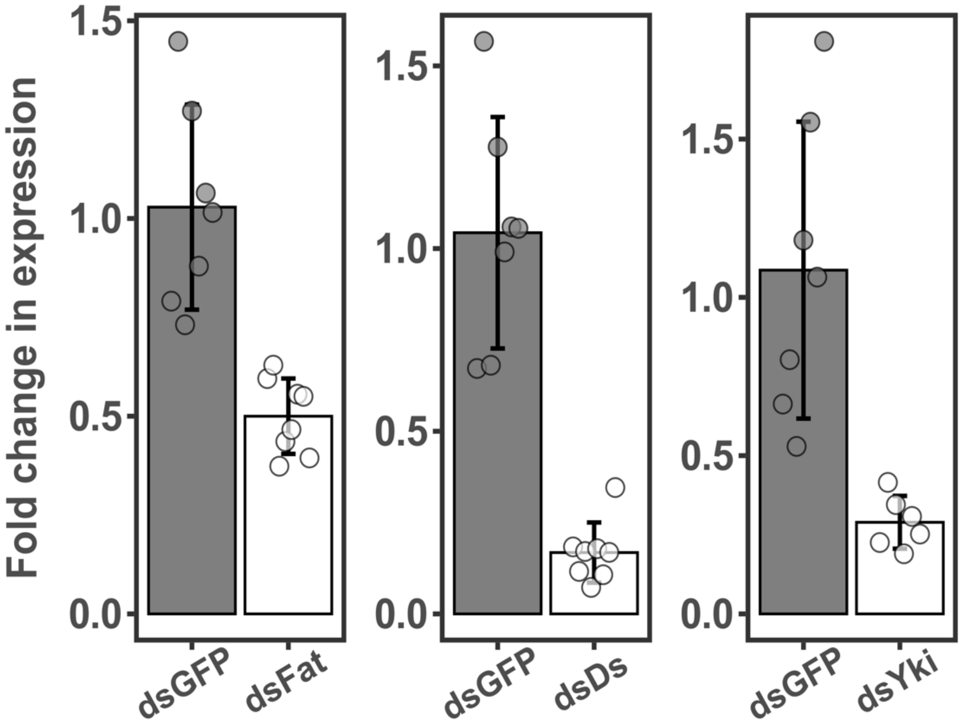

**Figure S7. RT-qPCR validation of the knockdowns.** The fold change in expression levels between the control (dsGFP) and the RNAi knockdowns is shown for the knockdowns that caused a significant change in wing morphology.

### Supplementary Results

#### *Genome assembly and annotation*

The new assembly considerably increased contiguity in comparison with the original I5K assembly. The number of scaffolds was reduced to half and both N50 and N80 increased by about 5×, but the process did not result in a chromosome-length scaffold assembly. The new assembly had a total length of 1,001,615,004 bp across N = 9,479 scaffolds. The longest scaffold was 13,987,836 bp, N50 = 2,219,597 bp (n = 121 scaffolds) and N80 = 696,899 bp (n = 365 scaffolds). The I5K *G. buenoi* 1.1 assembly, on the other hand, had a total length of 994,443,371 bp across N = 18,844 scaffolds, a longest scaffold of 2,551,663 bp, and with N50 = 412,278 (n = 674 scaffolds) and N80 = 138,063 bp (n = 1,908 scaffolds) (Poelchau *et al.*, 2015; Armisén *et al.*, 2018).

De novo repeat discovery with RepeatModeler (Flynn *et al.*, 2020) produced a library containing 146 novel LTR families and 3277 unclassified repeat families. The result of combining this library with Hemiptera repeats using RepeatMasker (Smit, Hubley and Green, 2013) resulted in a total of 28.7% of the scaffolded assembly being softmasked. Overall, the total masked bases included 15.1% retroelements (6.6% LINEs and 8.5% LTR retroelements), 11.6% DNA transposons (including 7.7% Tc1-IS630-Pogo) and 62.9% of interspersed repeats. Low-complexity repeats totaled 10.4% of the total masked length.

The initial consensus annotation determined from the softmasked assembly included 20,353 gene models and 22,232 subordinate transcript models. Eighty-nine percent of the manually curated protein sequences (n = 1,241) resulted in a single hit in the polished genome, and 1.9% yielded no hit. A total of 1,214 additional gene and transcript models were constructed from these alignments, which included high-quality duplicate gene models as described above. These models were incorporated into the annotation after removing overlapping consensus gene models. The final annotation included 20,431 gene models and 22,132 subordinate transcripts. Of the proteins encoded by these transcripts, 12,051 matched a FlyBase protein, 17,404 matched a protein from the *G. buenoi* 1.1 annotation, and 18,812 (92.1%) matched an OrthoDB 10v1 Hemiptera ortholog.

**References**

Armisén, D. *et al.* (2018) 'The genome of the water strider Gerris buenoi reveals expansions of gene repertoires associated with adaptations to life on the water', *BMC Genomics*, 19(1), p. 832. Available at: https://doi.org/10.1186/s12864-018-5163-2.

Flynn, J.M. *et al.* (2020) 'RepeatModeler2 for automated genomic discovery of transposable element families', *Proceedings of the National Academy of Sciences*, 117(17), pp. 9451–9457. Available at: https://doi.org/10.1073/pnas.1921046117.

Poelchau, M. *et al.* (2015) 'The i5k Workspace@NAL—enabling genomic data access, visualization and curation of arthropod genomes', *Nucleic Acids Research*, 43(D1), pp. D714–D719. Available at: https://doi.org/10.1093/nar/gku983.

Smit, A., Hubley, R. and Green, P. (2013) 'RepeatMasker Open-4.0'. Available at: https://www.repeatmasker.org (Accessed: 13 December 2022).
